# Mutations in SKI in Shprintzen-Goldberg syndrome lead to attenuated TGF-β responses through SKI stabilization

**DOI:** 10.1101/2020.10.07.330407

**Authors:** Ilaria Gori, Roger George, Andrew G. Purkiss, Stephanie Strohbuecker, Rebecca A. Randall, Roksana Ogrodowicz, Virginie Carmignac, Laurence Faivre, Dhira Joshi, Svend Kjaer, Caroline S. Hill

## Abstract

Shprintzen-Goldberg syndrome (SGS) is a multisystemic connective tissue disorder, with considerable clinical overlap with Marfan and Loeys-Dietz syndromes. These syndromes have commonly been associated with enhanced TGF-β signaling. In SGS patients, heterozygous point mutations have been mapped to the transcriptional corepressor SKI, which is a negative regulator of TGF-β signaling that is rapidly degraded upon ligand stimulation. The molecular consequences of these mutations, however, are not understood. Here we use a combination of structural biology, genome editing and biochemistry to show that SGS mutations in SKI abolish its binding to phosphorylated SMAD2 and SMAD3. This results in stabilization of SKI and consequently attenuation of TGF-β responses, in both knockin cells expressing an SGS mutation, and in fibroblasts from SGS patients. Thus, we reveal that SGS is associated with an attenuation of TGF-β-induced transcriptional responses, and not enhancement, which has important implications for other Marfan-related syndromes.

## INTRODUCTION

Shprintzen-Goldberg syndrome (SGS) is a multi systemic connective tissue disorder. Common features observed in SGS patients include craniofacial, skeletal and cardiovascular anomalies, ranging from heart valve defects to thoracic aortic aneurysms, all of which are also characteristic of Marfan syndrome (MFS) and Loeys-Dietz syndrome (LDS) (Cook et al., 2015a, Verstraeten et al., 2016, Loeys et al., 2005, Williams et al., 2007). In addition, SGS patients present with craniosynostosis, intellectual disability and skeletal muscle hypotonia (Shprintzen and Goldberg, 1982, Glesby and Pyeritz, 1989, Greally et al., 1998). All three syndromes have been linked to deregulation of the Transforming Growth Factor β (TGF-β) signaling pathway (Cannaerts et al., 2015).

The TGF-β family of ligands comprises the TGF-βs themselves, Activins, Nodal, bone morphogenetic proteins (BMPs) and growth differentiation factors (GDFs), and they play pleiotropic roles in embryonic development and tissue homeostasis. In addition, their signaling is deregulated in diverse pathologies (Miller and Hill, 2016). They exert their action by binding to type I and type II serine/threonine kinase receptors at the cell surface (TGFBR1 and TGFBR2 respectively, for the TGF-βs) (Massague, 2012). In the resulting ligand-bound heterotetrameric receptor complex, the type II receptor phosphorylates and activates the type I receptor, which in turn phosphorylates the intracellular mediators, the receptor-regulated SMADs (R-SMADs). Once phosphorylated, the R-SMADs (SMAD2 and SMAD3 in the case of TGF-β, Activin and Nodal) associate with the common mediator of the pathway, SMAD4. The resulting heterotrimeric complexes accumulate in the nucleus where they interact with other transcriptional regulators to activate or repress target gene expression (Massague, 2012). Two highly-related co-repressors, SKI and SKIL (formerly known as SnoN) act as negative regulators in the pathway (Deheuninck and Luo, 2009); see below).

The role of deregulated TGF-β signaling in Marfan-related syndromes is controversial. MFS is caused by loss-of-function mutations in the extracellular matrix protein, Fibrillin 1 (FBN1) (Dietz et al., 1991). These mutations are thought to increase the bioavailability of TGF-β ligands, as FBN1 binds the latent form of the TGF-βs (Neptune et al., 2003, Kaartinen and Warburton, 2003). Supporting the idea that excessive TGF-β signaling contributes to the manifestations of MFS, a TGF-β neutralizing antibody significantly improved the lung phenotype in a mouse model of MFS (homozygous Fbn1^mgΔ^) (Neptune et al., 2003, Cannaerts et al., 2015), and reduced the occurrence of aortic aneurysms in the Fbn1^C1039G/+^ mouse model of MFS (Habashi et al., 2006). Contradicting these results, others have shown that the aortopathy in the Fbn1^C1039G/+^ mouse model is not mediated by excessive TGF-β signaling, and in fact is exacerbated by loss of TGF-β signaling in smooth muscle cells (Wei et al., 2017). Furthermore, TGF-β signaling protects against abdominal aortic aneurysms in angiotensin II-infused mice (Angelov et al., 2017). This controversy emphasizes the importance of understanding exactly how TGF-β signaling is impacted in MFS. Furthermore, the related syndrome LDS is caused by pathogenic mutations in several different components of the TGF-β pathway, TGFBR1, TGFBR2, SMAD2, SMAD3, and the ligands, TGFB2 and TGFB3. These mutations all cause missense amino acid substitutions that have been either verified *in vitro*, or are predicted to be loss of function, implying that LDS is caused by attenuated TGF-β signaling (Horbelt et al., 2010, Cardoso et al., 2012, Schepers et al., 2018). However, paradoxically, histological and biochemical studies of aortic tissue derived from LDS patients reveal an apparent high TGF-β signaling signature (van de Laar et al., 2012, Gallo et al., 2014, Lindsay et al., 2012). SGS is caused by mutations in SKI, and both SMAD-mediated and non-SMAD-mediated TGF-β signaling has been reported to be increased in primary dermal fibroblasts from SGS patients (Doyle et al., 2012).

The co-repressors SKI and SKIL play important roles in a number of different cellular processes including proliferation, differentiation, transformation and tumor progression (Bonnon and Atanasoski, 2012). They are dimeric proteins that interact with both phosphorylated SMAD2 and SMAD3 (PSMAD2 or PSMAD3) via short motifs at their N-terminus, and with SMAD4 via a SAND domain (named after Sp100, AIRE-1, NucP41/75, DEAF-1) in the middle of both proteins (Deheuninck and Luo, 2009). Between these two domains lies a Dachshund homology domain (DHD), which is thought to also be important for R-SMAD binding (Wilson et al., 2004, Ueki and Hayman, 2003). SKI and SKIL both contain a leucine zipper domain in their C-termini, through which they dimerize (Deheuninck and Luo, 2009). They are negative regulators of TGF-β/Activin signaling, with two distinct mechanisms of regulation having been proposed. In one model, SKI and SKIL bind with SMAD4 to SMAD binding elements (SBEs) of TGF-β/Activin target genes, and recruit corepressors such as NCOR1 or SIN3A (Tokitou et al., 1999, Nomura et al., 1999, Stroschein et al., 1999, Deheuninck and Luo, 2009). They thus maintain the transcription of these target genes suppressed in the absence of signal. Upon TGF-β/Activin signaling, SKI and SKIL are rapidly degraded by the E3 ubiquitin ligase, RNF111 (formerly known as Arkadia), a process that requires SKI/SKIL binding to PSMAD2 or PSMAD3 (Le Scolan et al., 2008, Levy et al., 2007, Nagano et al., 2007). This then allows the activated SMAD3–SMAD4 complexes to bind the exposed SBEs and activate target gene transcription (Levy et al., 2007, Stroschein et al., 1999). In the competing model, SKI and SKIL act as repressors of active signaling simply by binding to PSMAD2 or PSMAD3 and SMAD4 in such a way as to disrupt the activated PSMAD2/PSMAD3–SMAD4 complexes (Luo, 2004, Ueki and Hayman, 2003, Wu et al., 2002). The heterozygous missense mutations that cause SGS have been mapped in SKI to the N-terminal R-SMAD-binding domain, with some small deletions and point mutations also found in the DHD, which is also necessary for R-SMAD binding (Carmignac et al., 2012, Doyle et al., 2012, Schepers et al., 2015). Thus, depending on the mechanism whereby SKI inhibits TGF-β/Activin signaling, loss of the interaction with PSMAD2/PSMAD3 would be predicted to have opposite effects on signaling output. If the PSMAD2/PSMAD3 interaction is required for SKI degradation, its loss would inhibit TGF-β signaling. However, if SKI binding to PSMAD2/PSMAD3 disrupts active SMAD complexes, then its loss would promote TGF-β signaling.

Here we use a combination of genome editing, structural biology, biochemistry and analysis of patient samples to elucidate the molecular mechanism underlying SGS and to resolve the paradox surrounding the role of TGF-β signaling in Marfan-related syndromes. We first determine at the molecular level how SKI/SKIL function in the TGF-β/Activin signaling pathways, and show that an intact ternary phosphorylated R-SMAD–SMAD4 complex is required for ligand-induced SKI/SKIL degradation. We demonstrate that the SGS mutations in SKI abolish interaction with PSMAD2 and PSMAD3 and this results in an inability of SKI to be degraded in response to TGF-β/Activin signaling. We go on to show that SKI stabilization results in an attenuation of the TGF-β transcriptional response in both knockin HEK293T cells and in fibroblasts from SGS patients. Our work unequivocally establishes that SGS mutations lead to an attenuated TGF-β response, which has major implications for all the Marfan-related syndromes.

## RESULTS

### A PSMAD2/3–SMAD4 ternary complex is essential for TGF-β/Activin-induced degradation of SKI/SKIL

To understand the consequences of SKI mutations in SGS and to resolve the paradox surrounding the function of TGF-β signaling in Marfan-related syndromes, we first set out to determine exactly how SKI and SKIL act as negative regulators of TGF-β and Activin signaling. We and other have previously demonstrated that SKI and SKIL are rapidly degraded upon TGF-β/Activin stimulation by the E3 ubiquitin ligase RNF111, and this requires PSMAD2 or PSMAD3 (Le Scolan et al., 2008, Levy et al., 2007, Nagano et al., 2007). Knockdown experiments suggested that SMAD4 was not necessary (Levy et al., 2007), but we subsequently showed that tumor cells deleted for SMAD4 or containing mutations in SMAD4 that abolish interactions with activated R-SMADs, abrogated TGF-β-induced degradation of SKI/SKIL (Briones-Orta et al., 2013). Whether the requirement for SMAD4 was direct or indirect was not clear.

To define the role of SMAD4 in SKI/SKIL degradation, we used CRISPR/Cas9 technology to delete SMAD4 in transformed embryonic kidney cells HEK293T, which express both SKI and SKIL, and in the human keratinocyte cell line, HaCaT, which predominantly express SKIL (Levy et al., 2007) (Figure 1 - source data 1). In wild type (WT) cells TGF-β/Activin induced rapid SKI and SKIL degradation, compared to cells treated with the TGFBR1 inhibitor, SB-431542 (Inman et al., 2002) (Figure 1A, B). Deletion of SMAD4 in multiple clones of both cell types abolished ligand-induced SKI/SKIL degradation (Figure 1A, B). We validated these SMAD4-null cell lines by demonstrating that transient expression of SMAD4 could rescue TGF-β/Activin-induction of the SMAD3–SMAD4 reporter, CAGA_12_-Luciferase (Figure 1 Supplement 1A, B). Furthermore, we could show that loss of SMAD4 inhibited the ligand-induced expression of a number of endogenous TGF-β and BMP target genes (Figure 1 Supplement 1C). By knocking out SMAD2 or SMAD3 individually or together, we also confirmed that these R-SMADs are absolutely required for TGF-β/Activin-induced degradation of SKI and SKIL, and act redundantly (Figure 1C) (Figure 1 - source data 1). Thus, R-SMADs and SMAD4 are all essential for TGF-β/Activin-dependent SKI/SKIL degradation.

**Figure 1.**
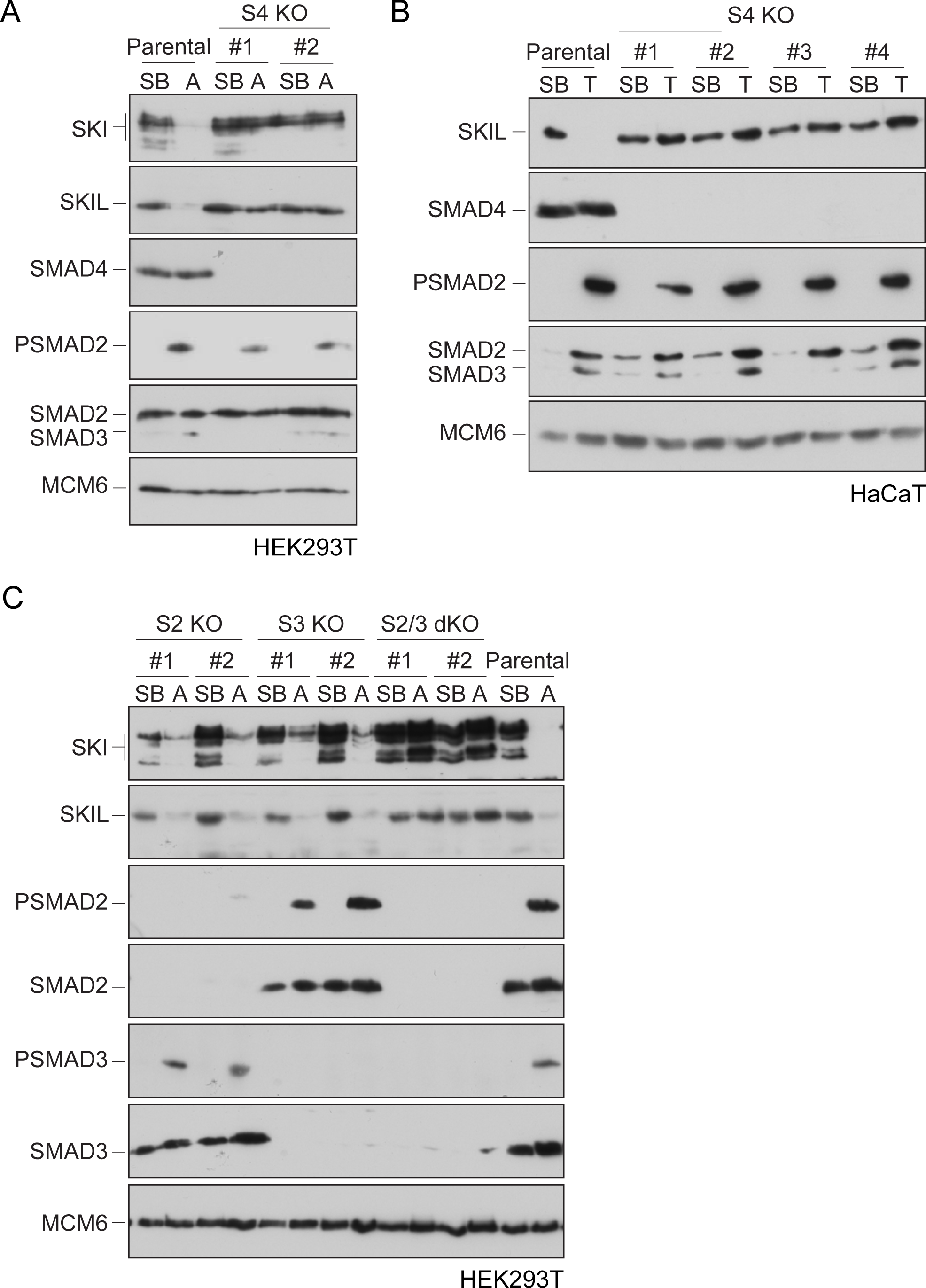
Requirement of SMAD2 or SMAD3 and SMAD4 for SKI and SKIL degradation. (A and C) The parental HEK293T cell line and two individual SMAD4 knockout clones (A) or two individual SMAD2, SMAD3 knockout clones, or two SMAD2 and SMAD3 double knockout clones (C) were incubated overnight with 10 μM SB-431542, washed out, then incubated with full media containing either SB-431542, or 20 ng/ml Activin A for 1 h, as indicated. Whole cells extracts were immunoblotted with the antibodies indicated. (B) Parental HaCaT and four individual SMAD4 knockout clones were treated as above, except that they were treated with 2 ng/ml TGF-β for 1 h instead of Activin A. Nuclear lysates were immunoblotted using the antibodies indicated. SB, SB431542; A, Activin A; T, TGF-β; S2, SMAD2; S3, SMAD3; S2/3, SMAD2 and SMAD3; S4, SMAD4; KO, knockout; dKO, double knockout.

In addition to forming a ternary complex with PSMAD2 or PSMAD3, SMAD4 has also been shown to interact directly with SKI and SKIL through their SAND domains (Wallden et al., 2017, Wu et al., 2002). To determine which of these SMAD4 interactions were important for TGF-β/Activin-induced SKI/SKIL degradation, we stably reintroduced enhanced GFP (EGFP) fusions of WT or mutated SMAD4 into HaCaT SMAD4-null cells. We selected two missense mutations on opposite faces of the C-terminal Mad homology 2 (MH2) domain of SMAD4: Asp351->His (D351H) and Asp537-->Tyr (D537Y) (Shi et al., 1997). These have been shown to occur naturally in the human colorectal cancer cell lines CACO-2 and SW948, and have lost the ability to bind phosphorylated R-SMADs (De Bosscher et al., 2004). In addition, we used the crystal structure of the MH2 domain of SMAD4 and the SAND domain of SKI, to design two mutations Ala433-->Glu (A433E) and Ile435-->Tyr (I435Y), that would be expected to abolish SMAD4 binding to SKI and SKIL (Wu et al., 2002).

We confirmed that these SMAD4 mutants behaved as expected in the rescue cell lines by testing their interaction with SKIL and R-SMADs by immunoprecipitation. As endogenous RNF111 triggers SKIL degradation in TGF-β/Activin dependent manner, the stable SMAD4-expressing HaCaT rescue cell lines were incubated with the proteasome inhibitor, MG-132 for 3 h prior TGF-β stimulation, to block SKIL degradation. As predicted, the D351H and D537Y SMAD4 mutants had lost their ability to bind SMAD2 upon TGF-β induction, but retained the interaction with SKIL. By contrast, A433E and I435Y SMAD4 mutants were unable to bind SKIL, but could interact with SMAD2 upon TGF-β stimulation (Figure 2A). Furthermore, as expected, D351H and D537Y SMAD4 mutants failed to rescue the ability of TGF-β to induce expression of CAGA_12_-Luciferase in HaCaT SMAD4-null cells or rescue TGF-β-induced transcription of target genes, but the A433E and I435Y SMAD4 mutants rescued these responses almost as well as WT SMAD4 (Figure 2 Supplement 1A, B).

**Figure 2.**
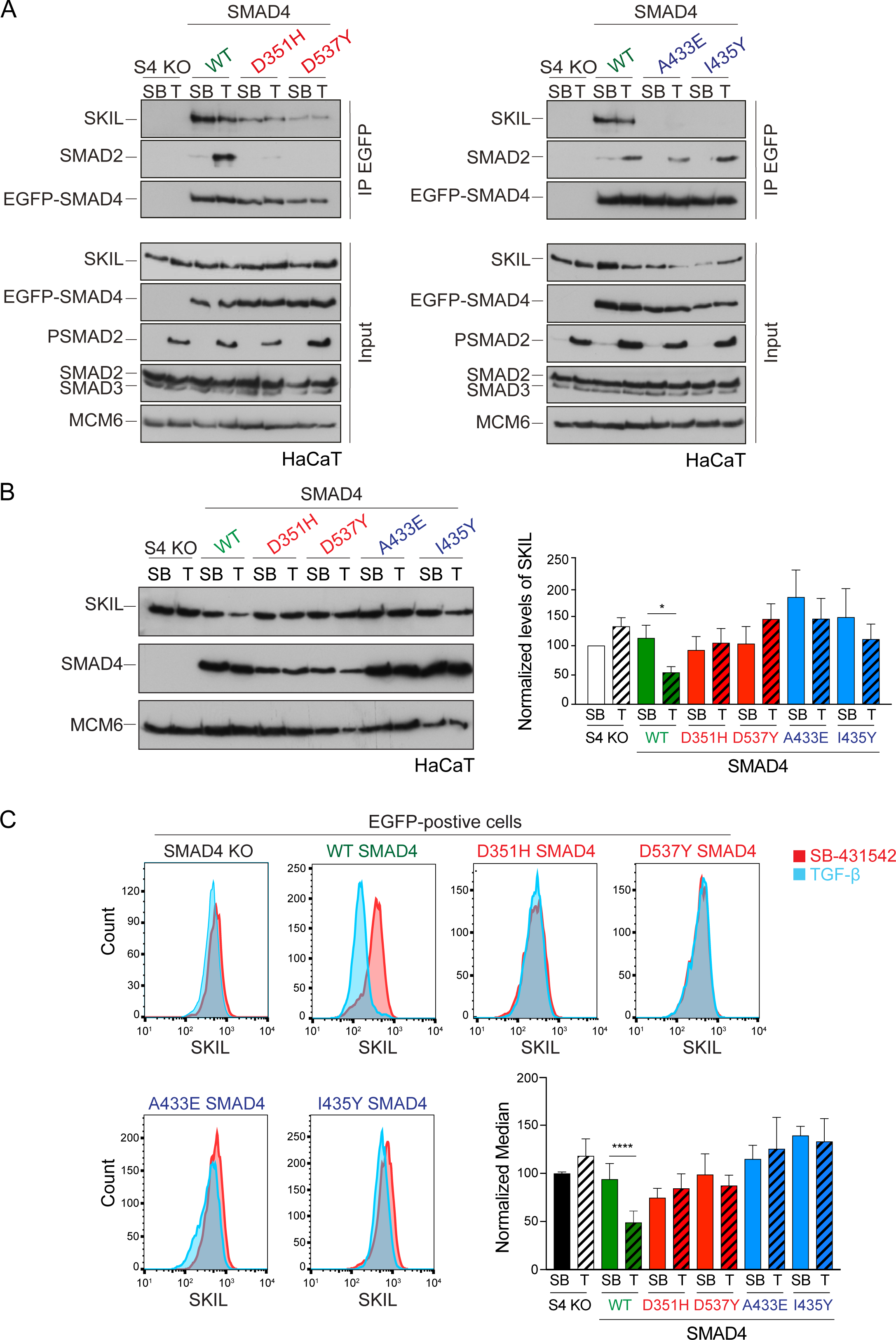
Characterization of the role of SMAD4 in TGF-β-induced SKIL degradation. (A– C) HaCaT SMAD4 knockout (S4 KO) cells were stably transfected with EGFP alone, or EGFP SMAD4 (WT) or with four different EGFP-SMAD4 mutants (D351H, D537Y, which abolish interaction with the RSMADs, and A433E and I435Y, which do not interact with SKIL). (A) Cells were incubated overnight with 10 µM SB-431542, washed out and pre-incubated with 25 μM MG-132 for 3 h, and then treated either with 10 μM SB-431542 or 2 ng/ml TGF-β for 1 h. Whole cell extracts were immunoprecipitated (IP) with GFP-trap agarose beads. The IPs were immunoblotted using the antibodies shown. Inputs are shown below. (B) Nuclear lysates were prepared from the HaCaT S4 KO cells stably transfected with EGFP alone or with EGFP-SMAD4 constructs as indicated, treated as in (A), but without the MG-132 step and immunoblotted using the antibodies shown. On the right the quantifications are the normalized average ± SEM of five independent experiments. The quantifications are expressed as fold changes relative to SB-431542-treated S4 KO cells. (C) Levels of SKIL in the EGFP-positive S4 KO rescue cell lines treated as in (B), assayed by flow cytometry. Each panel shows an overlay of the indicated treatment conditions. The red line indicates the SB-431542-treated sample, whereas the cyan line indicates the TGF-β-treated sample. Quantifications are shown bottom right. For each group, the percentage of the median fluorescence intensity normalized to the SB-431542-treated sample is quantified. Data are the mean ± SEM of five independent experiments. The *p* values are from one-way ANOVA with Sidak’s post hoc correction \**p*< 0.05; \*\*\*\**p*< 0.0001. SB, SB-431542; T, TGF-β.

Having demonstrated that these mutants behaved as designed, we asked which were able to mediate TGF-β-induced SKIL degradation, using three different assays. In a Western blot assay using nuclear extract, we found that reintroduction of WT SMAD4 in SMAD4-null cells caused a 50% reduction in SKIL levels in TGF-β-induced cells compared to those treated with SB-431542 (Figure 2B). However, none of the four SMAD4 mutants could rescue TGF-β-induced SKIL degradation (Figure 2B). We then established a flow cytometry assay to quantify SKIL protein stability in EGFP/EGFP-SMAD4-expressing cells (Figure 2C; Figure 2 Supplement 1C). Treatment with TGF-β for 1 h caused a 52% reduction in the relative median fluorescence intensity in the EGFP-SMAD4 WT-expressing cells, reflecting SKIL levels, compared to cells treated with SB-431542 (Figure 2C). However, for all four SMAD4 mutants tested, the median fluorescence was not decreased by TGF-β treatment (Figure 2C). Finally, we used an immunofluorescence analysis to monitor SKIL protein stability following TGF-β exposure. SMAD4-null cells showed strong nuclear staining of SKIL in the non-signaling condition (SB-431542), which remained unchanged by TGF-β treatment (Figure 3). Reintroduction of WT EGFP-SMAD4 conferred the ability to degrade SKIL upon TGF-β treatment, whereas none of the mutant SMAD4s were able to rescue SKIL degradation (Figure 3, arrows). Thus, all three assays demonstrate that a ternary R-SMAD–SMAD4 complex is absolutely necessary for TGF-β-induced SKIL degradation, as is the ability of SMAD4 to interact with SKIL itself. This suggests that within a canonical activated ternary SMAD complex, the R-SMAD component binds to the N-terminal region of SKIL/SKI, whilst SMAD4 binds the SAND domain, and both interactions are absolutely required for SKIL/SKI degradation.

**Figure 3.**
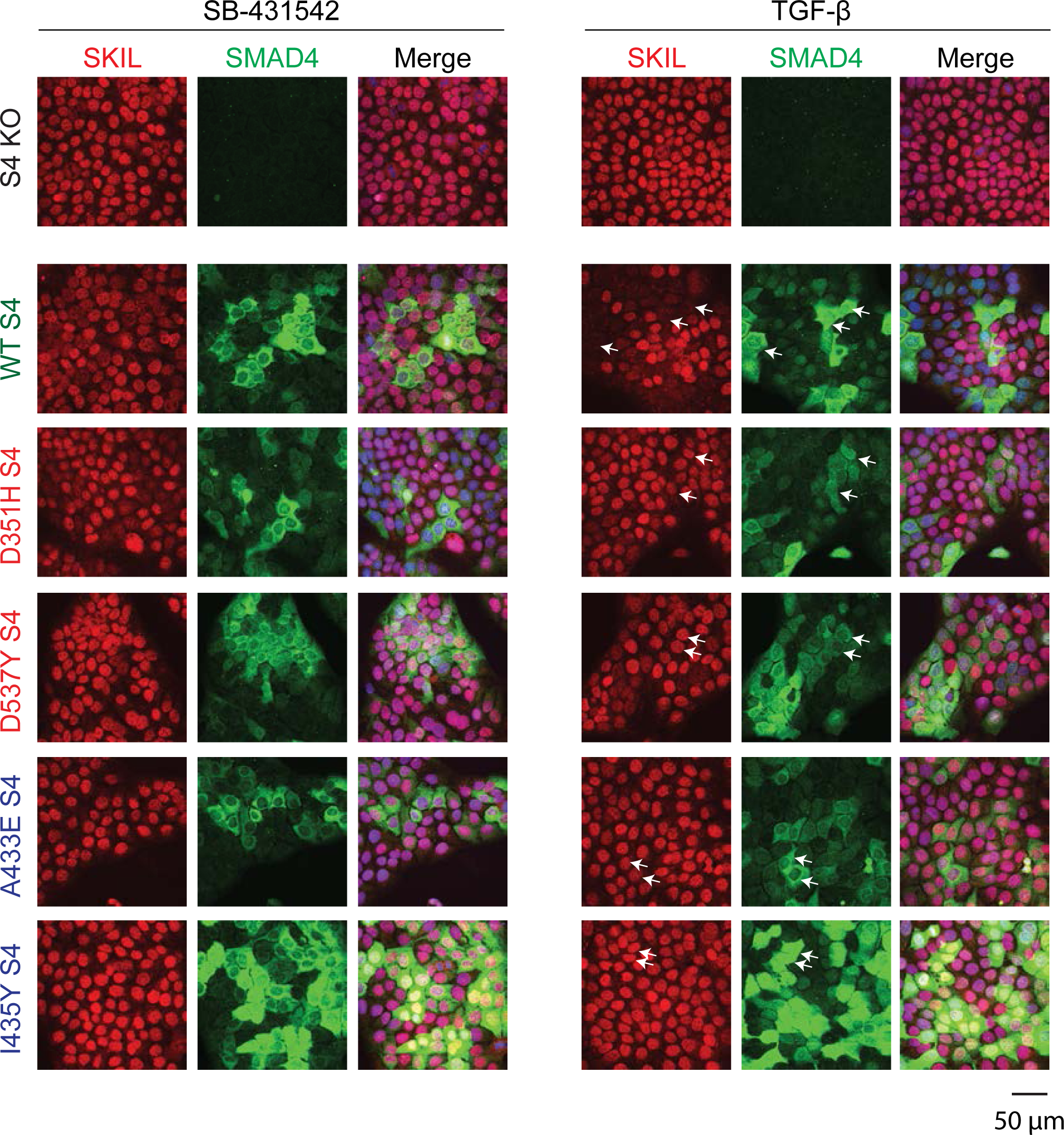
Visualization of TGF-β-induced SKIL degradation. HaCaT SMAD4 knockout (S4 KO) cells or those stably expressing EGFP SMAD4 WT or EGFP SMAD4 mutants were incubated overnight with 10 μM SB-431542, washed out and incubated for 1 h with 10 µM SB-431542 or with 2 ng/ml TGF-β. Cells were fixed and stained for EGFP (for SMAD4), SKIL and with DAPI (blue) to mark nuclei, and imaged by confocal microscopy. The merge combines SKIL, SMAD4 and DAPI staining. Arrows indicate examples of EGFP-expressing cells and corresponding levels of nuclear SKIL. Scale bar corresponds to 50 µm.

### SGS mutations inhibit the interaction of SKI with phosphorylated R-SMADs

We next investigated the consequences of the SGS mutations on SKI and SKIL’s ability to interact with the R-SMADs. SKI and SKIL share a highly conserved region at their N-terminus comprising the domain known to be important for R-SMAD binding (Deheuninck and Luo, 2009) (Figure 4 Supplement 1A). We first determined the minimal region of SKI required for R-SMAD binding using peptide pull down assays with biotinylated SKI peptides and whole cell extract from uninduced and TGF-β-treated HaCaT cells. This revealed that amino acids 11– 45 of SKI are sufficient for binding to PSMAD2 and PSMAD3 upon TGF-β stimulation, whilst the unphosphorylated SMADs did not bind to any of the SKI peptides (Figure 4 Supplement 1B). SMAD4 is also pulled down in these assays in a ligand-induced manner, by virtue of its interaction with the phosphorylated R-SMADs.

**Figure 4.**
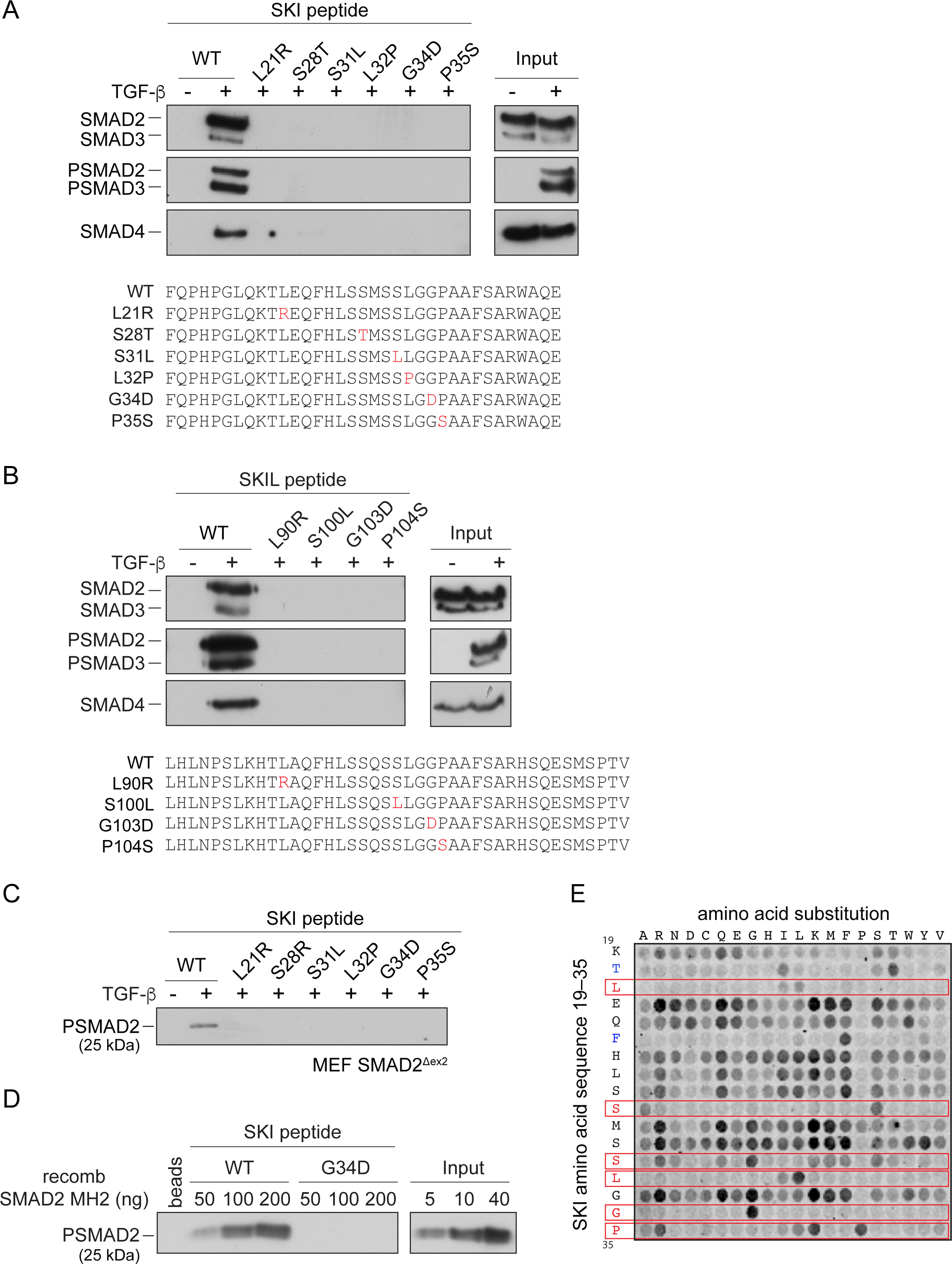
SGS mutations inhibit binding of SKI to SMAD2/3. (A and B) HaCaT cells were treated or not with 2 ng/ml TGF-β. A peptide pulldown assay was performed on whole cells extracts and pulldowns were immunoblotted with the antibodies indicated. Inputs are shown on the right. (A) Wild type (WT) SKI peptides corresponding to amino acids 11–45 or containing SGS point mutations as shown in red were used. (B) WT SKIL peptides corresponding amino acids 80–120 or containing mutations (in red) corresponding to SGS mutations in SKI were used. (C) WT SKI peptides or those containing SGS point mutations were used in pulldown assays with whole cell extracts of SMAD2-null mouse embryonic fibroblasts that express just the MH2 domain of SMAD2 (MEF SMAD2^Δex2^) (Das et al., 2009), treated with 2 ng/ml TGF-β. The untreated sample is only shown for the WT SKI peptide. A PSMAD2 immunoblot is shown. (D) A recombinant trimer of phosphorylated SMAD2 MH2 domain was used in a peptide pulldown assay with WT and G34D SKI peptides. A PSMAD2 immunoblot is shown, with inputs on the right. (E) Mutational peptide array of SKI peptides (amino acids 11–45), mutated at all residues between amino acids 19 and 35 was probed with a recombinant PSMAD3–SMAD4 complex, which was visualized using a SMAD2/3 antibody conjugated to Alexa 488. On each row the indicated amino acid is substituted for every other amino acid.

The SGS mutations discovered so far mostly cluster within this 11–45 region of SKI and a few deletions and point mutations have additionally been mapped in the DHD domain (Carmignac et al., 2012, Doyle et al., 2012, Schepers et al., 2015). The residues mutated are completely conserved, both between species, and also in the related protein, SKIL (Figure 4 Supplement 1A) (Carmignac et al., 2012). To determine the effect of these mutations on R-SMAD interaction, we introduced six different SGS mutations into the SKI peptide 11–45, and showed that they all prevented binding of PSMAD2 and PSMAD3, and as a result, also SMAD4 (Figure 4A). These results were also confirmed with the equivalent mutations in SKIL (Figure 4B). We proved that the interaction with SMAD2 was mediated via its MH2 domain using a mouse embryonic fibroblast cell line that expresses a truncated SMAD2 protein comprising just the MH2 domain (Piek et al., 2001, Das et al., 2009) (Figure 4C). We confirmed this using recombinant human phosphorylated SMAD2 MH2 domain produced in insect cells by co-expressing the SMAD2 MH2 domain with the kinase domain of TGFBR1 (Figure 4D). In both cases, the SGS mutations prevented interaction of the SKI peptide with the SMAD2 MH2 domain.

We next used a peptide array to gain a better understanding of which amino acids can be tolerated at the positions found to be mutated in SGS and to determine which other amino acids in this region of SKI are essential for the R-SMAD interaction. The SKI peptide corresponding to amino acids 11–45 was synthesized as an array on a cellulose sheet such that each residue in the sequence between residues 19 and 35 was substituted with all 19 alternative amino acids (Figure 4E). The array was probed with a recombinant PSMAD3– SMAD4 trimer, generated by co-expressing the SMAD3 and SMAD4 with the TGFBR1 kinase domain in insect cells. The PSMAD3–SMAD4 complex was then detected using a fluorescently-labeled SMAD2/3 antibody. Eight residues are intolerant to almost any amino acid substitution (Thr20, Leu21, Phe24, Ser28, Ser31, Leu32, Gly34 and Pro35). Strikingly, six of these residues are the amino acids known to be mutated in SGS patients, and the array results readily explain why these residues are mutated to a number of different amino acids in SGS (Figure 4E). In addition, Thr20 and Phe24 are also crucial residues for binding the PSMAD3–SMAD4 complex, but have not yet been reported as disease mutations. Mutations in the other nine amino acids do not impair the binding, and almost any other amino acid apart from proline can be tolerated at these positions.

### Crystal structure of the SKI peptide with the phosphorylated SMAD2 MH2 domain

To discover why these eight amino acids were so crucial for R-SMAD binding, and also to understand why SKI and SKIL only recognize phosphorylated R-SMADs, we solved the crystal structure of the SKI peptide (amino acids 11–45) with a phosphorylated homotrimer of the SMAD2 MH2 domain. We confirmed using SEC-MALLS that the phosphorylated SMAD2 MH2 domain was indeed trimeric in solution (Figure 5 Supplement 1A). Analysis of the binding affinity of the SKI peptide to the SMAD2 MH2 domain trimer indicated that the dissociation constant (*K*_d_) was in the low nanomolar range (Figure 5 Supplement 1B). The structure was determined by molecular replacement and refined at 2 Å resolution and readily explained why the crucial amino acids identified in the peptide were required for SMAD2 binding (Figure 5; Figure 5 Supplement 1C). The SKI peptide binds on the outside face of the MH2 domain at the so-called three helix bundle, comprising helices 3, 4 and 5 (Wu et al., 2001) (Figure 5A). The N-terminal helix of SKI packs against helix 3 of SMAD2, and the C-terminal portion of the SKI peptide, which contains the critical Gly34 and Pro35, forms a sharp turn that is stabilized by pi-stacking coordination between Phe24 of SKI, Trp448 of SMAD2 and Pro35 of SKI (Figure 5B). Moreover, the NE1 of the Trp448 side chain forms a H-bond to the main chain carbonyl group of Gly33, which in turn positions Pro35 for the interaction with Trp448 (Figure 5B). Furthermore, Glu270 in SMAD2 provides a pocket, which has a negatively charged base that ties down SKI Gly34 through hydrogen-bonding to its main chain amides. Other key interactions involving amino acids identified above as crucial for binding include the main chain carbonyl of Ser31, which forms a hydrogen bond to the ND1 of Asn387 in helix 3 (Figure 5C), and the hydroxyl group of SKI Thr20, which forms a hydrogen bond with the Gln455 at the end of helix 5 of SMAD2, and is nearly completely buried in the interface (Figure 5D). The two leucine residues (Leu21 and Leu32) that are mutated in SGS, are both buried in the structure (Figure 5E, F).

**Figure 5.**
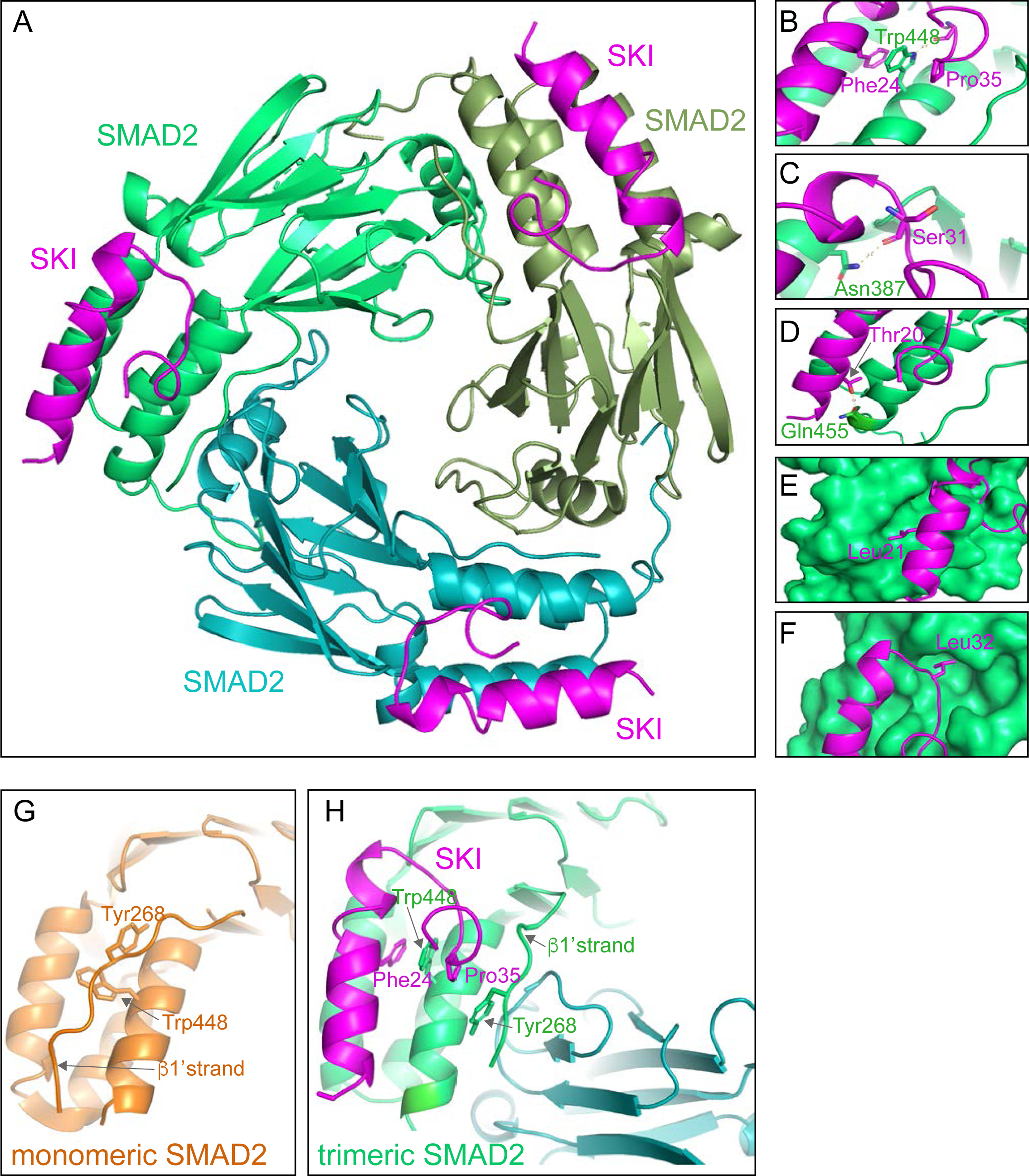
Crystal structure of PSMAD2 MH2 domain and N-terminal SKI peptide. (A) Crystal structure of the phosphorylated SMAD2 MH2 domain trimer (the three monomers are shown in bright green, cyan and olive) with the N-terminal SKI peptide amino acids 11–45 (purple). A ribbon representation is shown. (B–F) Close ups on key residues for SKI binding. SKI residues are shown in pink, and SMAD2 residues are in green. In B–D a ribbon representation is shown. In E and F, SMAD2 is shown as a surface representation and SKI as a ribbon. (G) A detail from the structure of monomeric SMAD2 MH2 domain with a peptide from ZFYVE9 (formerly called SARA) (Wu et al., 2000). Note that the β1’ strand that contains Tyr268 is locked in a hydrophobic pocket, forcing Trp448 into flattened orientation, incompatible with SKI binding. (H) A detail from the structure in (A) indicating how SMAD2 complex formation shifts the position of the β1’ strand and more particularly, Tyr268, allowing Trp448 to flip 90°, enabling it to stack with SKI residues Phe24 and Pro35.

Having demonstrated how SKI binds to SMAD2, we asked why SKI only binds phosphorylated R-SMADs and not monomeric unphosphorylated R-SMADs. To do this we compared the structure of the unphosphorylated SMAD2 MH2 domain bound to a region of ZFYVE9 (formerly called SARA) (Wu et al., 2000) with our current structure. It was clear that in the unphosphorylated SMAD2 structure, Tyr268 in the so-called β1’ strand (amino acids 261–274) is locked in a stable conformation in a hydrophobic pocket, and also forms a number of hydrogen bonds (Figure 5G). Crucially, this conformation forces Trp448 into flattened orientation, which is incompatible with SKI binding through the pi-stacking involving SKI Phe24, SMAD2 Trp448 and SKI Pro35 (Figure 5G). MH2 domain trimerization generates a new binding site for the β1’ strand on the adjacent MH2 domain subunit (Figure 5H; Supplementary Video 1). The central residue driving this is Tyr268. In the trimer the hydroxy group of Tyr268 makes hydrogen bond contact with the carbonyl group of Asp450 and the main chain of Lys451 on the adjacent MH2 domain subunit. As a consequence, Trp448 moves into an upright position in the trimer, allowing engagement with SKI. Thus, SKI can only bind SMAD2 in its phosphorylated trimeric state.

### Knockin of an SGS mutation into HEK293T cells inhibits Activin-induced SKI degradation and attenuates PSMAD3–SMAD4-mediated transcriptional activity

We have shown that the presence of SGS mutations prevent the interaction of SKI/SKIL with phosphorylated R-SMADs and have demonstrated that SKI/SKIL degradation requires an activated R-SMAD–SMAD4 complex. We therefore went on to investigate the functional effect of the SGS mutations on SKI degradation and TGF-β/Activin-induced transcriptional responses. To do this, we chose to focus on Pro35 because of its crucial role in forming the stacking interaction with Trp448 in SMAD2. In SGS patients Pro35 is mutated to Ser or Gln (Carmignac et al., 2012, Doyle et al., 2012, Schepers et al., 2015), substitutions not tolerated in activated R-SMAD–SMAD4 binding (Figure 4E).

We used CRISPR/Cas9 technology with a single-stranded template oligonucleotide to knock in the Pro35 → Ser (P35S) mutation into HEK293T cells, and we efficiently generated a number of homozygous clones (Figure 6 Supplement 1A). In three independent clonal cell lines carrying the P35S SKI mutation, the binding to endogenous phosphorylated SMAD2 was severely compromised, compared with WT SKI (Figure 6A). The binding to SMAD4, however, was unchanged in the mutant cell lines, as the SGS mutations do not affect the SKI SAND domain which is responsible for SMAD4 binding (Figure 6A). To assess the impact of the P35S SKI mutation on the SKI and SKIL degradation, cells were treated with Activin for 1 or 2 h and SKI/SKIL levels determined by immunoblotting. At both time points, we clearly demonstrated that P35S SKI levels remained stable, whilst in the parental cell lines SKI protein level is almost entirely degraded after 1 h of Activin treatment (Figure 6B). The presence of mutated SKI had no effect on the Activin-induced degradation of SKIL in these lines (Figure 6B). Thus, the P35S mutation renders SKI completely resistant to ligand-induced degradation.

**Figure 6.**
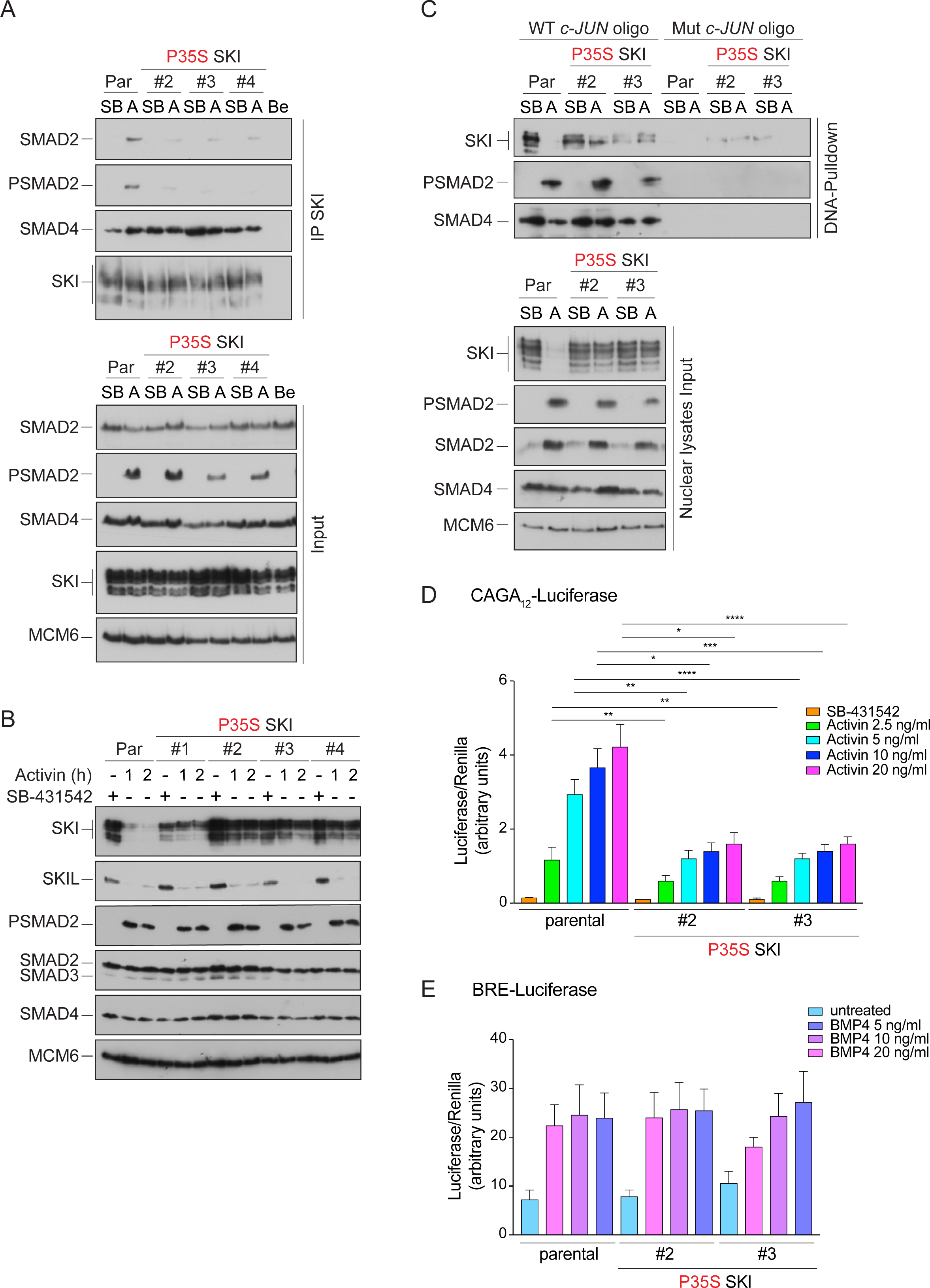
Knockin of an SGS mutation into SKI in HEK293T cells inhibits SKI degradation and inhibits Activin-induced transcription. (A) Parental HEK293T and three independent P35S SKI knockin clones were incubated overnight with 10 μM SB-431542, washed out and treated for 3 h with 25 μM MG-132 and then with either SB-431542 or 20 ng/ml Activin A for an additional 1 hr. Whole cell lysates were immunoprecipitated (IP) with SKI antibody or beads alone (Be). The IPs were immunoblotted using the antibodies shown. Inputs are shown below. (B) Parental HEK293T and four independent P35S SKI knockin clones were incubated with 10 μM SB-431542 overnight, washed out and incubated with either SB-431542 or 20 ng/ml Activin for the times indicated. Whole cell lysates were immunoblotted using the antibodies indicated. (C) Cell were treated as in (B), and nuclear lysates were prepared and analyzed by DNA pull-down assay using the wild type c-Jun SBE oligonucleotide or a version mutated at the SMAD3–SMAD4 binding sites (top panel). Inputs are shown in the bottom panel. HEK293T parental and two independent P35S SKI knockin clones stably transfected with the CAGA_12_-Luciferase reporters (D) or the BRE-Luciferase reporter (E) and TK-Renilla as an internal control. Cells were serum starved with media containing 0.5% FBS and 10 μM SB-431542 overnight. Subsequently, cells were washed and treated with Activin A (D) or BMP4 (E) at the concentrations indicated for 8 h. Cell lysates were prepared and assayed for Luciferase and Renilla activity. Plotted are the means and SEM of seven (D) or four (E) independent experiments, with the ratio of Luciferase:Renilla shown. *, *p*<0.05; **, *p*<0.01; ***, *p*<0.001; ****, *p*<0.000. The *p*-values are from two-way ANOVA with Tukey’s post-hoc test. A, Activin; SB, SB-431542; Par, parental.

To determine if SGS mutations had the same effect in SKIL, we introduced a G103V mutation into SKIL, corresponding to the SGS mutation G34V in SKI (referred to as SKIL ΔS2/3). Transfection of G103V SKIL in HEK293T cells led to reduction of SMAD2 binding in parental cells (Figure 6 Supplement 1B). The residual binding was mediated via SMAD2’s interaction with SMAD4, as it was lost in the SMAD4 knockout cells (Figure 6 Supplement 1B). Binding of SMAD4 in absence or presence of signal was unaffected by the mutation. As observed above for SKI, the SGS mutation in SKIL led to resistance to Activin-induced degradation (Figure 6 Supplement 1C), indicating that the R-SMAD interaction was essential. In addition, we made a version of SKIL with mutations in the SAND domain (R314A, T315A, H317A and W318E) that rendered it unable to interact with SMAD4 (referred to as SKIL ΔS4). This mutant was also not degraded upon Activin stimulation (Figure 6 Supplement 1C), demonstrating an essential requirement for SMAD4 binding.

SKI and SKIL bind DNA in conjunction with SMAD4 at SBEs of TGF-β/Activin target genes in the absence of ligand stimulation. The ligand-induced degradation of SKI and SKIL then allows the activated R-SMAD–SMAD4 complexes access to the SBEs to activate transcription of target genes (Levy et al., 2007, Stroschein et al., 1999). We hypothesized that if the SGS mutations render SKI resistant to ligand-induced degradation, then mutant SKI and SMAD4 would remain bound to the DNA. To test this, we used a DNA pulldown assay with an oligonucleotide corresponding to SBEs from the *JUN* promoter (Levy et al., 2007). Consistent with our prediction, both WT and P35S SKI bound the SBEs with SMAD4 in the absence of signal, but after Activin stimulation, the binding of WT SKI is lost, whilst the binding of P35S SKI is retained (Figure 6C).

The SKI–SMAD4 complex bound to SBEs in the absence of signal are transcriptionally repressive (Levy et al., 2007, Stroschein et al., 1999). Since P35S SKI remains bound with SMAD4 in Activin-stimulated cells, we reasoned that this would inhibit Activin-induced gene expression. To address this, we stably expressed luciferase reporters (CAGA_12-_Luciferase and BRE-Luciferase), together with TK-Renilla as an internal control (Dennler et al., 1998, Korchynskyi and ten Dijke, 2002), in parental HEK293T cells and in two independent clones of the knockin P35S SKI cells. The CAGA_12_-Luciferase reporter responds to TGF-β and Activin, is induced by PSMAD3–SMAD4 complexes and is sensitive to SKI and SKIL levels, whilst the BRE-Luciferase reporter is induced by SMAD1/5–SMAD4 complexes in response to BMPs and is not affected by SKI and SKIL (Levy et al., 2007). Strikingly, we found a significant reduction in Activin-induced CAGA_12-_Luciferase activity in the P35S SKI cells compared to parental cell line (Figure 6D), while BMP4-induced BRE-Luciferase activity was similar in all cell lines (Figure 6E). The results indicate that SGS mutations in SKI lead to inhibition of TGF-β/Activin-induced transcription mediated by PSMAD3–SMAD4 complexes.

### Dermal fibroblasts from SGS patients exhibit an attenuated transcriptional TGF-β response

To gain further insights into the functional consequences of the SGS mutations we obtained dermal fibroblasts from two SGS patients, a female carrying the heterozygous point mutation L32V, and a male carrying a heterozygous deletion of 12 base pairs corresponding to codons 94–97 (ΔS94-97) (Carmignac et al., 2012). Both patients present the classical features of SGS such as marfanoid habitus and intellectual disability, whilst the patient carrying L32V mutation also manifests craniosynostosis. In addition, we obtained dermal fibroblast from a healthy male subject as a control.

We investigated whether the SGS mutations rendered SKI resistant to TGF-β-induced degradation in the dermal fibroblasts, as demonstrated in the knockin HEK293T cells. Indeed, in control fibroblasts SKI expression is abrogated upon TGF-β stimulation after 1 h, while in the SGS-derived fibroblasts SKI protein remains relatively stable (Figure 7A). Note that these cells are heterozygous for the mutation, while the HEK293T P35S SKI knockins used above were homozygous, thus accounting for the incomplete SKI stabilization exhibited by the SGS fibroblasts, compared with the knockin cells.

**Figure 7.**
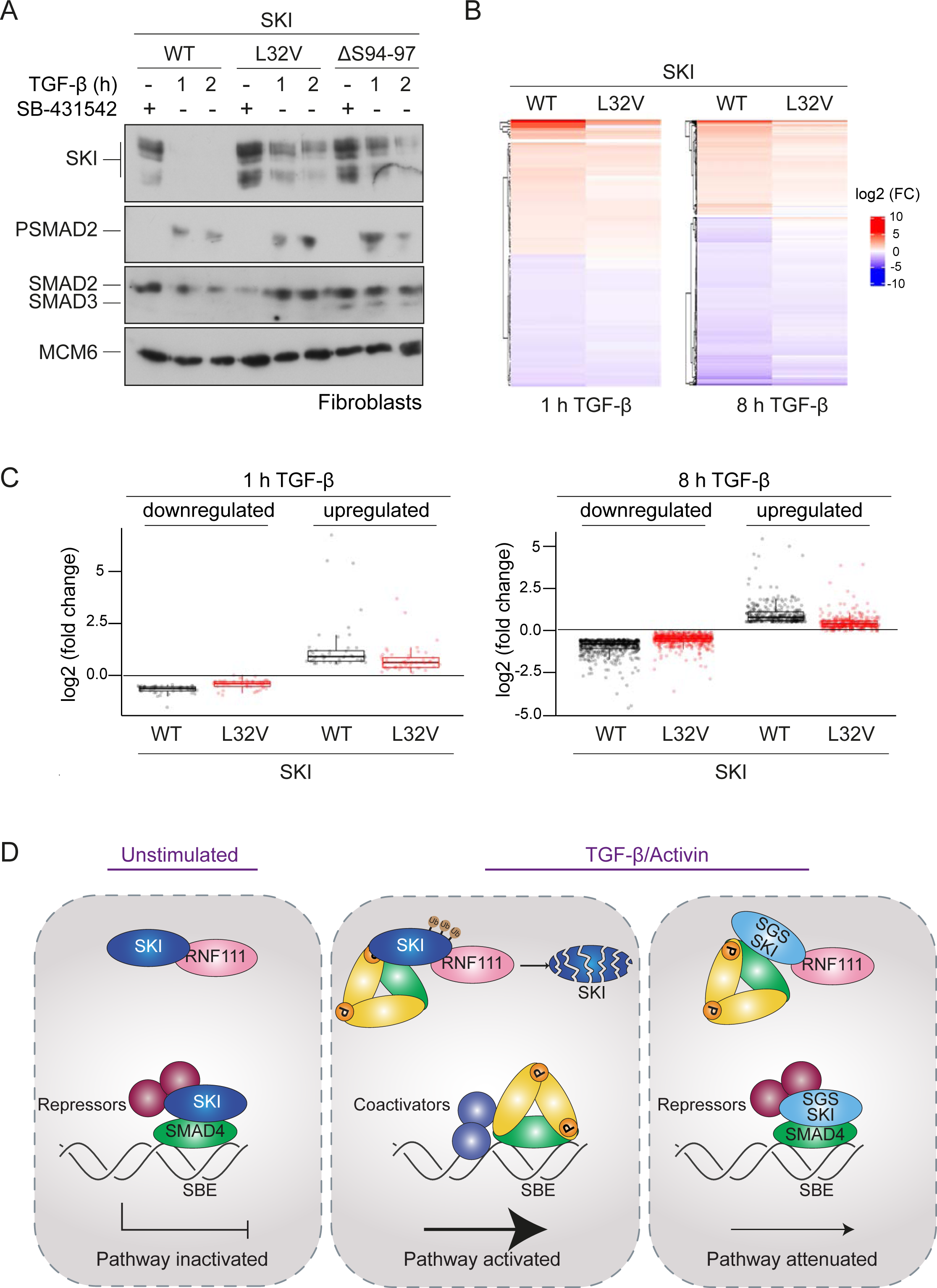
SGS mutations in SKI inhibit TGF-β-induced transcriptional responses in fibroblasts derived from SGS patients. (A) Fibroblasts derived from a healthy subject carrying WT SKI and from two SGS patients carrying the L32V or the ΔS94-97 heterozygous mutations in SKI were incubated overnight with 10 μM SB-431542, washed out and either re-incubated with SB-431542 or with 2 ng/ml TGF-β for the times indicated. Whole cell lysates were immunoblotted using the antibodies indicated. (B) Hierarchically-clustered heatmaps of log_2_FC values (relative to the SB-431542-treated samples) showing the expression of TGF-β-responsive genes in the healthy fibroblasts and the L32V SKI fibroblasts after 1 h and 8 h of TGF-β treatment, analyzed by RNA seq. Four biological replicates per condition were analyzed. The genes shown are those for which the TGF-β inductions were statistically significant in the healthy fibroblasts, but non-significant in the L32V fibroblasts. (C) The same data as in (B) are presented as box plots. (D) Model for the mechanism of action of WT SKI and mutated SKI. The left panel shows the unstimulated condition. In the nuclei, SKI (blue) is complexed with RNF111 (pink) and is also bound to DNA at SBEs with SMAD4 (green) forming a transcriptionally repressive complex with other transcriptional repressors (maroon). In the middle panel, TGF-β/Activin stimulation induces the formation of phosphorylated R-SMAD–SMAD4 complexes (yellow and green), which induce WT SKI degradation by RNF111. This allows an active PSMAD3–SMAD4 complex to bind SBEs and activate transcription. In the right panel, SGS-mutated SKI (light blue) is not degraded upon TGF-β/Activin stimulation, due to its inability to interact with PSMAD2 or PSMAD3. It therefore remains bound to SMAD4 on DNA, leading to attenuated transcriptional responses.

To determine the effect of SKI stabilization on global TGF-β-induced transcription we performed genome-wide RNA-sequencing (RNA-seq) in three different conditions: SB-431542-treated cells (non-signaling condition), 1 h TGF-β-treated and 8 h TGF-β-treated, and compared the L32V and ΔS94-97 SKI fibroblasts to control fibroblasts. The samples separated in a principal component analysis according to the cell line used and the treatment performed (Figure 7 Supplement 1A), and we confirmed that the differentially-enriched genes after TGF-β treatment in the control fibroblasts were characteristic of pathways related to TGF-β signaling (Figure 7 Supplement 1B). We then performed a pairwise comparison between the TGF-β-treated and SB-431542-treated samples for each of the cell lines individually. We found that of the 339 genes that were differentially expressed in normal fibroblasts after 1 h of TGF-β treatment, 60% (202 genes) were induced or repressed less efficiently in the L32V mutant fibroblasts, and of these, 97 genes were not significantly differentially expressed in the mutant cells at all (Figure 7 – source data 1, Figure 7B, C). After an 8 h TGF-β induction, we found that 4769 genes were differentially expressed in the normal fibroblasts, and of these 75% (3556 genes) were induced or repressed less efficiently in the L32V mutant cells, and 880 genes were not significantly differentially expressed in the mutant cells at all (Figure 7 – source data 1, Figure 7B, C). We observed similar results when comparing the TGF-β responses in the normal fibroblasts versus ΔS94-97 SKI fibroblasts, although the effects were less dramatic (Figure 7 – source data 1, Figure 7 Supplement 1C, D). To illustrate the magnitude of these effects we validated six gene expression profiles (*ISLR2, CALB2, SOX11, ITGB6, HEY1, COL7A1*) in the normal versus mutant fibroblasts by qPCR (Figure 7 Supplement 2).

Thus, we conclude that the presence of SGS point mutations in SKI that render it resistant to ligand-induced degradation, result in attenuated TGF-β responses for a substantial subset of target genes.

## DISCUSSION

### SGS mutations in SKI lead to stabilization of SKI and attenuated TGF-β/Activin transcriptional responses

In this study, we have resolved the mechanism of action of SKI and the related protein SKIL, which has allowed us to elucidate the molecular consequences of SKI mutations in SGS. Our proposed mechanism of how SKI acts as a transcriptional repressor of TGF-β/Activin signaling in health and disease is illustrated in Figure 7D, and our results suggest that the mechanism of action of SKIL is equivalent. In the absence of ligand stimulation, in both healthy and diseased cells, SKI and SKIL bind in conjunction with SMAD4 at SBEs of TGF-β/Activin target genes. Here, they repress transcription by recruiting corepressors such as NCOR1 or SIN3A (Tokitou et al., 1999, Nomura et al., 1999, Stroschein et al., 1999, Deheuninck and Luo, 2009). We and others have also shown that in unstimulated cells SKI and SKIL interact with the E3 ubiquitin ligase, RNF111, although this binding *per se* does not lead to SKI/SKIL degradation (Le Scolan et al., 2008, Levy et al., 2007, Nagano et al., 2007). In healthy cells, upon TGF-β/Activin stimulation, SKI/SKIL form a complex with a canonical PSMAD2/PSMAD3–SMAD4 trimer, which induces rapid degradation of SKI/SKIL via RNF111 (Le Scolan et al., 2008, Levy et al., 2007, Nagano et al., 2007). A possible mechanism explaining this would be that the binding of the PSMAD2/PSMAD3–SMAD4 complex induces an activating conformational change in RNF111, although this has not yet been demonstrated. Degradation of SKI and SKIL removes the repressors from the SBEs, allowing access of activated PSMAD3–SMAD4 complexes to the SBEs to regulate transcription of target genes. In SGS cells, mutated SKI can no longer interact with PSMAD2/PSMAD3, and it is therefore not degraded upon TGF-β/Activin signaling. It thus remains bound with SMAD4 to SBEs, resulting in an attenuation of TGF-β/Activin transcriptional responses. To demonstrate the functional consequences of the SGS mutations, we used genome-wide RNA-seq analysis of fibroblasts derived from SGS patients and have shown that SGS mutations indeed lead to a reduction in the magnitude of TGF-β transcriptional responses.

### The mechanism underlying SKI and SKIL function

Since the discovery that SKI and SKIL interact with SMAD2 and SMAD3, and act as negative regulators of TGF-β/Activin pathways (Luo et al., 1999, Stroschein et al., 1999, Sun et al., 1999), two different mechanisms of action have been proposed. One mechanism is as described in the paragraph above – an initial version of which was first proposed in 1999 (Stroschein et al., 1999). The second mechanism was based on the crystal structure of the SKI SAND domain with the MH2 domain of SMAD4 (Wu et al., 2002). In this crystal structure, the binding of SKI with SMAD4, which is mediated via the I-loop of the SKI SAND domain with the L3 loop of SMAD4, was mutually exclusive with the binding of SMAD4 to activated R-SMADs, which also requires the L3 loop of SMAD4. Thus, the authors concluded that the mechanism whereby SKI (and by analogy, SKIL) inhibited TGF-β/Activin signaling was by binding to the activated R-SMADs and SMAD4 in such a way as to disrupt the phosphorylated R-SMAD–SMAD4 complexes required for transcriptional activation (Wu et al., 2002). This mechanism has been supported by the observation that overexpression of SKI and SKIL inhibits TGF-β-induced functional responses (Luo, 2004). The two mechanisms are fundamentally different. In the first, SKI and SKIL are constitutive repressors that need to be degraded to allow pathway activation. In the second, SKI and SKIL act as inducible repressors, as they repress only upon ligand induction, by virtue of their ability to disrupt activated SMAD complexes.

Our biochemical analysis of the role of SMAD4 in SKI/SKIL function now resolves the controversy between the two models. Firstly, the efficient SKIL and SKI degradation that we and others have observed upon ligand stimulation in effect rules out the second model, at least in the first hours after ligand induction, as there would be little or no nuclear SKI or SKIL to disrupt activated SMAD complexes. Secondly, we clearly demonstrate for the first time that an intact functional phosphorylated R-SMAD–SMAD4 trimer is required to bind to SKIL to induce its ligand-dependent degradation. The key piece of evidence for this comes from our analysis of the inability of SMAD4 point mutants to restore ligand-induced SKIL degradation in SMAD4-null HaCaTs. Critically, we show that SMAD4 mutants that cannot form a canonical activated R-SMAD–SMAD4 trimer cannot rescue ligand-induced SKIL degradation, neither can SMAD4 mutants that do not interact with the SAND domain of SKIL. Thus, we conclude that TGF-β/Activin-induced SKIL degradation occurs only when SKIL interacts simultaneously with phosphorylated R-SMADs and with SMAD4, which in turn must interact with each other in a transcriptionally active trimer. This strongly indicates that SKIL binding to the R-SMAD– SMAD4 complex does not disrupt it.

This conclusion is supported by a recent crystal structure of the SAND domain of SKIL with the SMAD4 MH2 domain (Wallden et al., 2017). This structure revealed that SKIL interacts with SMAD4 in two states: an “open” and a “closed” conformation. In the open conformation the authors showed that SKIL can bind the R-SMAD–SMAD4 without intermolecular clashes or further structural readjustment, whereas in the closed state, structural reorganization within the SMAD heterotrimer is required to allow binding of SKIL, as has been observed in the previous structure of SKI with SMAD4 (Wu et al., 2002). Molecular modelling has subsequently confirmed that SKIL in open the conformation forms a stable ternary SKIL–SMAD3–SMAD4 complex (Ji et al., 2019). Furthermore, surface plasmon resonance indicated only one dominant binding mode for SKIL and SMAD4, leading to the conclusion that the open conformation is the biologically and functionally relevant mode, and that the closed conformation may be the result of crystal packing forces (Wallden et al., 2017). Note that the residues which allow the binding in the open conformation are highly conserved between SKI and SKIL, suggesting that both repressors bind to an intact activated R-SMAD– SMAD4 complex, which is required for their degradation. This would exclude the disruption model and thus favors the degradation model. It will now be important to solve the structure of an activated R-SMAD–SMAD4 trimer with the N-terminal half of SKI or SKIL that contains the R-SMAD binding motif, the DHD domain and the SAND domain that contacts the SMAD4 moiety. The role of the DHD domain is particularly intriguing as SGS mutations also occur in this domain (Carmignac et al., 2012, Doyle et al., 2012, Schepers et al., 2015) and appear from our patient sample analysis to have a similar effect on inhibiting ligand-induced SKI degradation.

Our structural data also elegantly explain why SKI and SKIL only bind to phosphorylated SMAD2 and SMAD3 in the context of an activated SMAD trimer, and not to monomeric SMAD2 or SMAD3. The key residue for this discrimination is Trp448 in SMAD2, the equivalent of which would be Trp406 in SMAD3. In the trimer, this residue is in a conformation compatible with stacking between Phe24 and Pro35 of SKI. In the monomer, however, it is rotated approximately 90°, prohibiting SKI binding. The binding mode of SKI to SMAD2 is distinct from that of other SMAD2-binding partners, for example, the transcription factor FOXH1, which contains two binding motifs. One of which, the so-called SIM binds SMAD2 in both monomeric and trimeric forms, whilst the so-called FM only binds phosphorylated trimeric SMAD2, because it recognizes the interface of the SMAD trimer (Miyazono et al., 2018, Randall et al., 2002, Randall et al., 2004). It will be interesting in the future to discover whether any other SMAD2/3-binding partner uses the same mode of interaction as SKI and SKIL.

### The molecular mechanism of SGS

As discussed in the Introduction, SGS is a Marfan-related syndrome, with patients exhibiting many of the same features characteristic of MFS and LDS. These syndromes have been considered as TGF-β signalopathies, as the causal mutations are either direct components of the TGF-β signaling pathway, or as in the case of MFS, a component of the microfibrils in the ECM, FBN1, that is known to bind latent TGF-β in complex with latent TGF-β binding proteins (LTBPs) (Cannaerts et al., 2015, Ramirez et al., 2004, Robertson et al., 2015). There has been controversy over whether the manifestations of these syndromes result from too little TGF-β signaling, or too much. This is obviously a crucial issue to resolve, as it is influencing the types of treatments being developed for patients with these syndromes.

The first suggestion that MFS resulted from excessive TGF-β came from mouse models, where key phenotypes could be rescued by a TGF-β neutralizing antibody (Habashi et al., 2006, Neptune et al., 2003). However, later studies using a potent murine anti-TGF-β antibody, or genetic methods for reducing TGF-β signaling have not corroborated these findings, and have worsened, rather than improved disease in MFS mouse models (Cook et al., 2015b, Holm et al., 2011, Lindsay et al., 2012, Wei et al., 2017). Furthermore, administration of small molecule inhibitors of the TGF-β type I receptor, or a pan-TGF-β neutralizing antibody, have been associated with serious adverse cardiovascular toxicities, such as valve defects, similar to those found in MFS (Anderton et al., 2011, Stauber et al., 2014, Mitra et al., 2020). The finding that mutations in FBN1 that prevent binding of LTBPs might result in lower levels of TGF-β signaling, rather than excessive signaling, is not so surprising given the more recent understanding of how TGF-β is activated. In order for mature TGF-β ligands to be released from the latent complex, either force has to be applied via integrins, to partially unfold the cleaved TGF-β pro-domain allowing release of the mature domain, or the pro-domain must be degraded by proteases (Dong et al., 2017, Rifkin et al., 2018, Robertson and Rifkin, 2016). For the traction mechanism to occur, the integrin must be anchored to the actin cytoskeleton, and the LTBP must be tethered to ECM, via FBN1 and fibronectin. Thus, release of latent TGF-β alone is not therefore sufficient to produce active TGF-β ligands.

Consistent with the view that lower levels of TGF-β signaling might be responsible for MFS, the mutations that give rise to LDS are all loss-of-function mutations in TGF-β pathway components (Schepers et al., 2018). Paradoxically though, signatures of higher TGF-β signaling were observed over time in mouse models of LDS (Gallo et al., 2014, MacFarlane et al., 2019). However, in this case, the pathology could not be rescued by neutralizing TGF-β activity. One possibility with both the mouse models of MFS and LDS is that the mutations do initially lead to lowered TGF-β signaling, but over time cells compensate by up-regulating either TGF-β ligands themselves, or other TGF-β family ligands that signal through PSMAD2/PSMAD3, ultimately leading to the enhanced signaling signatures observed.

With respect to SGS, there has been much less research into the consequences of the SGS mutations on TGF-β signaling responses. Based on the SMAD complex disruption model of SKI/SKIL action, it has been assumed that loss-of-function mutations in a negative regulator would lead to an increase in TGF-β signaling, and in fact this has been used to support the idea that the Marfan-related syndromes are caused by excessive TGF-β signaling (Doyle et al., 2012, Gallo et al., 2014). Here we unequivocally show that the opposite is true. We demonstrate that these mutations lead to loss of ligand-induced SKI degradation. As a result, the stabilized SKI remains bound to SBEs with SMAD4 as a repressive complex, and hence, a subset of TGF-β/Activin transcriptional responses are attenuated. We have proven this in both HEK293T knockin cells and in patient-derived fibroblasts. Moreover, we also find no evidence for increased PSMAD2 or PSMAD3 signaling. Indeed, neither model of SKI function would actually predict that the SKI mutations would affect levels of phosphorylated R-SMADs, since SKI acts downstream of R-SMAD phosphorylation. Finally, our finding that SGS mutations in SKI lead to its stabilization and are not equivalent to loss of SKI function, also explains why patients with 1p36 deletion syndrome, who are haploinsufficient for SKI, do not have SGS. However, unsurprisingly, many of the same organs are affected in both syndromes (Colmenares et al., 2002, Zhu et al., 2013).

It will now be important to use animal models to explore how attenuation of transcription of specific TGF-β/Activin target genes leads to the manifestations of SGS, and to understand why SGS patients exhibit additional defects compared with LDS and MFS patients. We anticipate that our new understanding that the SKI mutations lead to attenuation of TGF-β responses will resolve the paradoxes surrounding the role of aberrant TGF-β signaling in the other Marfan-related disorders, and will help inform the development of new therapeutic approaches.

## METHODS

### Cell lines

HEK293T and HaCaT cells were obtained from the Francis Crick Institute Cell Services and cultured in Dulbecco’s modified Eagle’s medium (DMEM) supplemented with 10% fetal bovine serum (FBS) and 1% Penicillin/Streptomycin (Pen/Strep). All CRISPR-Cas9 edited cell lines were cultured in the same media. Dermal fibroblasts from healthy subjects were kindly provided by David Abraham (UCL-Medical School Royal Free Campus). L32V and ΔS94-97 SKI dermal fibroblasts were obtained from Laurence Faivre and Virginie Carmignac (Université de Bourgogne UMR1231 GAD, Dijon, France) under the ethics of the GAD collection, number DC2011-1332 (Carmignac et al., 2012). The mutations were confirmed by Sanger Sequencing and RNA sequencing. The fibroblasts were all cultured in DMEM supplemented with 10% FBS, 1% Pen/Strep and 1% Insulin Transferrin Selenium (ITS, Thermo Fisher). Mouse embryo-derived fibroblasts harboring the homozygous null allele *Smad2*^*ex2*^ (MEF SMAD2^ex2^) (Piek et al., 2001) were maintained in DMEM supplemented with 10% FBS and 1% Pen/Strep. All cell lines have been banked by the Francis Crick Institute Cell Services, and certified negative for mycoplasma. The identity of all cell lines was also authenticated by confirming that their responses to ligands and their phenotype were consistent with published history. All the cell lines are listed in the Key Resource Table.

### Ligands, chemicals and cell treatments

Ligands and inhibitors were used at the following concentrations: TGF-β (PeproTech), 2 ng/ml; Activin A (PeproTech), 20ng/ml; BMP4 (PeproTech), 20 ng/ml; SB-431542 (Tocris), 10 µM; MG-132 (Tocris), 25 µM. All treatments were performed in full serum or where required, in serum-starved (0.5% FBS) DMEM. Unless otherwise stated, cells were incubated with 10 µM SB-431542 overnight to inhibit autocrine signalling, then were washed three times with warm media and stimulated with either Activin A or TGF-β. For proteasome inhibition, cells were treated for 3 h with 25 µM MG-132 prior stimulation with Activin A or TGF-β.

### Plasmids

Plasmids are listed in the Key Resource Table. CAGA_12_-Luciferase, BRE-Luciferase, TK-Renilla, pEGFP-C1, pEGFP-SMAD4 were as described previously (Dennler et al., 1998, Korchynskyi and ten Dijke, 2002, Levy et al., 2007, Nicolas et al., 2004). The pEGFP-SMAD4 mutants (D351H and D537Y) were generated by swapping the mutated SMAD4 coding region from the EF-HA vector into pEGFP-C1 (De Bosscher et al., 2004). The pEGFP-SMAD4 SMAD4 mutants (A433E and I435Y) were generated by PCR using oligonucleotides listed in the Key Resource Table. EF-Flag-SKIL G103V was generated from pEF-FLAG-SKIL by PCR using oligonucleotides listed in the Key Resource Table, while the mutant containing the R314A, T315A, H317A and W318E mutations was generated by synthesizing the SKIL region between BSTEII and AVRII sites containing the mutations and cloning that fragment into pEF-FLAG-SKIL. For plasmids used for generating recombinant proteins, see below.

### Transfections, generation of stable cell lines and reporter assays

Cells were transfected with the appropriate plasmids using Fugene 6 (Roche) according to the manufacturer’s instructions. Luciferase reporter assays were performed as previously described, using Dual-Glo assay system (Promega) following the manufactures’ instructions (Levy et al., 2007).

HaCaT SMAD4 KO lines stably expressing either EGFP or EGFP-SMAD4 WT or mutants were generated by transfecting the cells with the appropriate plasmids. Transfected cells were selected with 500 µg/ml of G418 (Invitrogen), then FACS-sorted for EGFP-positive cells and expanded. EGFP expression was confirmed by microscopy. To generate stable HEK293T cell lines expressing either CAGA_12_-Luciferase or BRE-Luciferase together with TK-Renilla, cells were transfected with the appropriate plasmids together with a plasmid carrying the puromycin resistance gene (pSUPER-retro-puro; OligoEngine). Cells were then selected with 2 μg/ml puromycin (Sigma).

### CRISPR/Cas9-mediated knockout of SMADs in HEK293T and HaCaT cells

For the generation of knockin or knockout HEK293T cells, a parental clone was selected which was a representative clone from the HEK293T pool that showed a robust Activin-induced SKI degradation and responded to TGF-β family ligands in the same way as the starting pool. For HaCaTs, a pool of cells was used as starting material for knockouts.

For SMAD2 and SMAD3 knockouts, a guide RNA in the MH2 domain of the protein was selected, whereas for SMAD4, two guide RNAs were picked, one targeting the MH1 domain and the other targeting at the end of the MH2 domain. The guide RNAs are shown in the Key Resource Table. The guides RNAs were expressed from the plasmid pSpCas9(BB)-2A-GFP (pX458) (Addgene, #48138) (Ran et al., 2013). HEK293T and HaCaT were transfected with the appropriate plasmid and for the double knockout SMAD2 and SMAD3, the two plasmids were transfected simultaneously. 48 h after transfection, cells were sorted for EGFP expression, plated as single cells in 96-well plates and screened by Western blot to assess the loss of the protein. For HEK293Ts, two knockout clones for SMAD2, SMAD3, SMAD2/SMAD3 and SMAD4 were used in these studies. For HaCaTs, four independent SMAD4 knockout clones were used. The sequences of the knockout alleles are shown in Figure 1 – source data 1.

### Knockin of P35S SKI at the endogenous locus

To introduce the P35S mutation into SKI, a gRNA was selected immediately downstream of codon 35. A 120-bp ssODN, where the codon CCG (P35) was mutated to TCC (S35) and codons 33 and 34 where silently mutated from GGC to GGA, was made and purified by Sigma (Key Resource Table). The ssODN contained phosphorothioate bonds between the first two and last two nucleotides at the 5’ and 3’ ends respectively, to avoid ssODN degradation by endogenous nucleases. The silent mutations at codons 33 and 34 were introduced to increase the specificity of the downstream screening primer. The mutation at codon 35 also disrupts the PAM sequence.

Cells were cotransfected with the pX458 plasmid expressing the gRNA and 10 μM ssODN using Fugene 6. After 48 h, cells were FACS-sorted for GFP expression, and plated as single cells in 96-well plates. Subsequently, clones were consolidated and from replicate plates, genomic DNA was extracted using Quickextract DNA extraction solution (Lucigen) according to manufacturer’s instructions. PCR was performed using a universal reverse primer and two different forward primers: a primer which allows detection of WT SKI and one which detects the P35S SKI (Key Resource Table). Clones positive for P35S SKI knockin were selected, verified by Sanger Sequencing and used for further analysis.

### Western blotting, immunoprecipitations, DNA pulldowns and immunofluorescence

Whole-cell extracts were prepared as previously described (Inman et al., 2002), while nuclear lysates were prepared according to (Wong et al., 1999). Western blots were carried out using standard methods. The list of the antibodies used is shown in the Key Resource Table. Immunoprecipitations using GFP-Trap beads (Cromotek) were performed according to the manufacturer’s instructions. Immunoprecipitations using antibodies coupled to protein G-Sepharose beads (Sigma) were as described previously (Levy et al., 2007).

DNA pulldown assays were performed as previously described with some modifications (Levy et al., 2007). Nuclear lysates were extracted using buffer containing 360 mM NaCl, and the DNA pulldowns were performed in the presence of a 40 μg of non-biotinylated mutant oligonucleotide to reduce non-specific binding. The oligonucleotides corresponding to WT and mutated SBE of the *JUN* promoter are shown in the Key Resource Table. Immunofluorescence was performed as previously described (Pierreux et al., 2000), except that cells were washed and fixed for 5 min in methanol at -20 C (Levy et al., 2007). Nuclei were counter stained with DAPI (0.1 *µ*/ml). Imaging was performed on a Zeiss Upright 780 confocal microscope. Z-stacks were acquired for all channels and maximum intensity projection images are shown.

For all of these techniques, a representative experiment of at least two biological repeats is shown.

### Flow cytometry

After 1 h of TGF-β induction HaCaT SMAD4 KO rescue cells, expressing either EGFP or EGFP-SMAD4 fusions were washed, trypsinized and pelleted. Cell pellets were fixed with methanol for 5 min at -20 C. Fixed cells were incubated with primary antibody against SKIL. Cells were then washed three times in phosphate buffered saline (PBS) and incubated with secondary antibody conjugated with anti-rabbit Alexa 647. As a negative control, we used cells incubated with secondary antibody only. Antibodies used are listed in the Key Resource Table. Subsequently, cells were washed three times with PBS, and pellets were resuspended with 500 µl of PBS and filtered to achieve a single cell suspension. Cells were then analyzed for EGFP fluorescence on an LSRII flow cytometer (BD Biosciences), gated for viable, single cells. We then quantified the fluorescence emitted by Alexa 647 in cells expressing EGFP as a measure of SKIL protein. The FlowJo program was used to analyze the results.

### Peptide pulldown assays and peptide array

For peptide pulldowns, N-terminal biotinylated peptides were synthesized by the Peptide Chemistry Facility at the Francis Crick Institute using standard procedures. The peptide pulldown assays were performed as described previously (Randall et al., 2002). Where recombinant protein was used, it was dissolved in buffer Y (50 mM Tris-HCl, pH 7.5, 150 mM NaCl, 1 mM EDTA 1% (v/v) NP-40), and used at the concentrations given in the legend to Figure 4. Peptides sequences are given in the Key Resource Table.

A peptide array was generated using peptides corresponding to SKI amino acids 11-45, of which the amino acids 19–35 were mutated one by one to every other amino acid. Arrays were synthesised on an Intavis ResPepSL Automated Peptide Synthesiser (Intavis Bioanalytical Instruments, Germany) on a cellulose membrane by cycles of N(a)-Fmoc amino acids coupling via activation of carboxylic acid groups with diisopropylcarbodiimide (DIC) in the presence of hydroxybenzotriazole (HOBt), followed by removal of the temporary α-amino protecting group by piperidine treatment. Subsequent to chain assembly, side chain protection groups are removed by treatment of membranes with a deprotection cocktail (20 ml 95% trifluoroacetic acid, 3% triisopropylsilane, 2% water 4 h at room temperature) then washing (4 x dichloromethane, 4 x ethanol, 2 x water, 1 x ethanol), prior to being air dried.

Peptide array membranes were blocked with 5% Milk in 0.01% Tween 20 in PBS and then incubated with purified PSMAD3–SMAD4 complex (see below) overnight. Subsequently, the membranes were washed and incubated with an antibody against SMAD2/3 (BD Biosciences) conjugated to Alexa 488 using the Zenon® Mouse IgG Labeling Kits (Life Technology) according to the manufacturer’s instructions. Fluorescence was detected where binding occurred between SKI and PSMAD3–SMAD4 complex and measured with a Typhoon FLA 9500 biomolecular imager (GE Healthcare).

In all peptide experiments, a representative of at least two biological repeats is shown.

### Generation of phosphorylated SMAD2 MH2 domain in insect cells

The cDNA encoding a fusion protein consisting of GST followed by a 3C cleavage site and the human SMAD2 MH2 domain (residues 241–465) was inserted into the MCSI of the pFastBac™ Dual vector (Thermo Fisher Scientific) in the Sal1 and Spe1 restriction sites. The cDNA encoding a constitutively-active version of the human TGFBR1 kinase domain (residues 175–503 with a T204D mutation) was cloned into the MCSII using the SmaI and SphI restriction sites. A high-titre baculovirus (>10^8^ pfu/ml) was generated using standard published protocols (Fitzgerald et al., 2006). Expression of the trimeric phosphorylated SMAD2 MH2 domain was performed by infecting Sf21 cells at a density of 1.5×10^6^ cells/ml at MOI:1 and incubating for 72 h at 27°C with rotation at 110 r.p.m. Cells were harvested by centrifugation at 1000×g for 10 min and stored at -80°C until required.

### Expression of the human phosphorylated SMAD3–SMAD4 complex

An expression construct was generated encoding GST fused full length human SMAD3 (with 3C protease site) and inserted into the MCSI of the pFastBac™ dual. As with the SMAD2 MH2 domain construct, the constitutively-active human TGFBR1 was cloned into MCSII. Another vector encoding full length human SMAD4 alone was also constructed and inserted into the pBacPAK plasmid. The resulting vectors were used to generate high titre virus (>10^8^ pfu/ml) using a standard published protocol. For expression of the phosphorylated SMAD3–SMAD4 complex both viruses were used to infect cultures of Sf21 insect cells at a density of 1.5×10^6^ cells/ml at a MOI:1. Infected cultures were allowed to grow for 72 h at 27°C with rotation at 110 r.p.m. Cells were harvested by centrifugation at 1000×g for 10 minutes and stored at - 80°C until required.

### Purification of trimeric phosphorylated SMAD2 MH2 domain and phosphorylated SMAD3–SMAD4 complexes

The same procedure was used to purify both phosphorylated SMAD2 MH2 domain trimers and phosphorylated SMAD3–SMAD4 complexes. Typically, 500 ml of infected Sf21 cells were lysed in 30 ml of a lysis buffer consisting of 50 mM HEPES (pH 8.0), 250 mM NaCl, 10% (v/v) glycerol, 1% (v/v) Triton X-100, 10 mM β-glycerophosphate, 1 mM NaF, 10 mM benzamidine and 1 mM DTT supplemented with 5 µl BaseMuncher (2500 U/µl), 5 mM MgCl_2_ and phosphatase (phosphatase inhibitor cocktail 3, Sigma) and protease inhibitors (EDTA free cOmplete™ protease inhibitors, Roche). After incubation for 20 min, the suspension was sonicated to ensure complete lysis. The insoluble fraction was pelleted by centrifugation (100,000×g at 4°C for 30 min). The soluble fraction was incubated with 500 µl bed volume of Glutathione 4B Sepharose (Cytiva) for 2 h at 4°C with gentle agitation. The resin was washed extensively with buffer containing 50 mM HEPES (pH 7.5), 200 mM NaCl, 1 mM DTT. The phosphorylated SMAD2 MH2 domain or phosphorylated SMAD3–SMAD4 complexes were eluted from the GSH resin by cleavage with GST-3C protease (20 µg) in 5 ml wash buffer overnight at 4°C with gentle agitation. The proteins were concentrated to 0.5 ml and applied to a S200 10/300 Increase (Cytiva) size exclusion column equilibrated with 50 mM HEPES (pH 7.5), 200 mM NaCl, 5% (v/v) glycerol, 1 mM DTT. Fractions were analyzed by SDS-PAGE and concentrated to 2 mg/ml and snap frozen as 50 µl aliquots and stored at -80°C until required. Purified phosphorylated SMAD2 MH2 domain was quantified using a molar extinction coefficient value of 39,420 M^-1^cm^-1^. The molar extinction coefficients used for SMAD3 and SMAD4 were 623,370 and 70,820 M^-1^cm^-1^ respectively.

### Size exclusion chromatography with multiangle laser light scattering (SEC-MALLS)

The trimeric arrangement of the phosphorylated SMAD2 MH2 domain was confirmed by SEC-MALLS. In brief, an S200 10/300 Increase column was attached to an AKTA Micro FPLC system (GE Healthcare), which was connected to a Heleos Dawn 8+ followed by an Optilab TRex (Wyatt). For data collection, 100 µl of a 2 mg/ml stock of phosphorylated SMAD2 MH2 domain was injected and data collected at 0.5 ml/min for 60 min. The data were analyzed by ASTRA6.1 software.

### Biolayer interferometry

Biolayer interferometry was carried out using an Octet RED96 instrument (ForteBio). Biotinylated N-SKI peptide (residues 11–45) was immobilized on streptavidin-coated biosensors (ForteBio) at a concentration of 1 μg/ml in buffer containing 50 mM HEPES pH 7.5, 200 mM NaCl, 1 mg/ml BSA and 0.1% Tween-20) for 100 s. The immobilization typically reached a response level of 2 nm. Association and dissociation curves were obtained through addition of a dilution series of trimeric phosphorylated SMAD2 MH2 domain complex (15.6 to 1.95 μM) for 100 s followed by dissociation in buffer for 350 s using the Octet acquisition software. The binding data were fitted using the Octet analysis software.

### Crystallization of the N-SKI–SMAD2 MH2 domain complex

The SKI peptide (amino acids 11–45) was synthesized by the Peptide Chemistry Group and added in a 2:1 ratio to the phosphorylated SMAD2 MH2 domain trimer. The complex was concentrated to 6 mg/ml and subject to crystallization trials. Initial screening gave rise to fine needles, in several conditions, and these were used as seedstock for rescreening. This gave rise to 25 µm crystals, with a cubic morphology, in the Crystal Screen Cryo screen (Hampton Research); the condition being 1.5 M Ammonium sulphate, 0.15 M K Na Tartrate, 0.08 M Na_3_ Citrate and 25% (v/v) glycerol. The crystals were flash-frozen in liquid nitrogen and data were collected on the I24 beamline at Diamond Light Source (DLS). Data was processed automatically using the DLS Xia2/XDS pipeline (Winter, 2010, Kabsch, 2010). The crystals belong to the I2_1_3 space group and diffracted to a resolution of 2.0 Å.

### Structure determination

Molecular replacement was undertaken with the CCP4 program Phaser (McCoy et al., 2007), utilising pdbfile 1khx (with the C-terminal tail removed), as the search model (Winn et al., 2011). Initial structure refinement was undertaken with Refmac (Murshudov et al., 2011), with manual model building in Coot (Emsley et al., 2010), before switching to Phenix.Refine to finalize the model (Liebschner et al., 2019, Afonine et al., 2012). Coordinates and data are available from the Protein Data Bank, with accession code: 6ZVQ.

### RNA extraction, qRT-PCR and RNA-sequencing

Total RNA was extracted using Trizol (Thermo Fisher Scientific) according to the manufacturer’s instructions. cDNA synthesis and qPCRs were performed as described (Gronroos et al., 2012). Primer sequences are listed in the Key Resource Table. All qPCRs were performed with the PowerUp SYBR Green Master Mix (Thermo Fisher Scientific) with 300 nM of each primer and 2 µl of diluted cDNA. Fluorescence acquisition was performed on either a 7500 FAST machine or QuantStudio 12 Flex (Thermo Fisher Scientific). Calculations were performed using the ΔΔCt method, and levels of mRNA are expressed as fold change relative to untreated or SB-431542-treated control cells. Means ± SEM from at least three independent experiments are shown. Results were analyzed using GraphPad Prism 8 software and statistics were performed on these data using one way or two-way Anova as stated in the Figure legends.

For the RNA sequencing experiment four biological replicates were used. Total RNA was extracted, and the quality of the RNA was assessed using a bioanalyzer (Agilent). Libraries were prepared using the KAPA mRNA HyperPrep kit (Roche) and single end reads were generated using an Illumina HiSeq 4000.

### RNA sequencing analysis

Raw reads were quality and adapter trimmed using cutadapt-1.9.1 (Martin, 2011) prior to alignment. Reads were then aligned and quantified using RSEM-1.3.0/STAR-2.5.2 (Dobin et al., 2013, Li and Dewey, 2011) against the human genome GRCh38 and annotation release 89, both from Ensembl. TPM (Transcripts Per Kilobase Million) values were also generated using RSEM/STAR. Differential gene expression analysis was performed in R-3.6.1 (Team, 2009) using the DESeq2 package (version 1.24.0) (Love et al., 2014). Differential genes were selected using a 0.05 false-discovery rate (FDR) threshold using the pairwise comparisons between time points within each cell line individually. Normalization and variance-stabilizing transformation (VST) was applied on raw counts before performing principal component analysis and euclidean distance-based clustering.

For the heatmaps, we selected those genes that were significantly differentially expressed in the control cell line with an absolute log2 (fold change) of at least 0.5, not significantly differentially expressed in the SKI-mutated cell lines, and detected with at least a TPM value of 2 in the control cell lines in any condition. We then visualized the log2 (fold change) for each time point against the control condition. Heatmaps were generated in R-3.6.1 using the ComplexHeatmap (Gu et al., 2016) package (version 2.0.0). The raw data files have been uploaded to GEO (GSE155292).

### Quantifications

Quantification of Western blots was performed by densitometry measurements of each lane using ImageJ software. The measurements were normalized to the loading control in the same blot. In each case, quantifications were normalized to the SB-431542-treated samples. Quantification for the flow cytometry was performed by measurement of the fluorescence emitted by Alexa 647 in EGFP expressing single cells as a measure of SKIL protein, and levels were normalized to the SB-431542-treated sample in the SMAD4 knockout cells. The program FlowJo was used to analyze the results.

### Statistical analysis

Statistical analysis was performed in Prism 8 (GraphPad). At least three independent experiments were performed for statistical analysis unless otherwise specified in the figure legends. Normalized values were log transformed for the statistical analysis. For comparison between more than two groups with one variable one-way analysis of variance (Anova) was used followed by the Sidak’s correction test. For comparison between groups that have been split on two independent variables, two-way analysis of variance (Anova) was performed followed by Tukey’s multiple comparison tests.

## ACKNOWLEDGMENTS

We would like to thank David Abraham for the normal dermal fibroblasts. We are very grateful to the Francis Crick Institute Advanced Sequencing Facility, the Light Microscopy Facility, the Flow Cytometry Facility, Cell Services, the Genomics Equipment Park and to the Peptide Chemistry Facility. We thank all the members of the Hill lab and Davide Coda, Daniel Miller, and Anassuya Ramachandran for helpful discussions and very useful comments on the manuscript. This work was supported by the Francis Crick Institute which receives its core funding from Cancer Research UK (FC001095), the UK Medical Research Council (FC001095), and the Wellcome Trust (FC001095).

The authors declare no competing interests.

## AUTHOR CONTRIBUTIONS

I.G. and C.S.H designed the project and experimental approaches. I.G. performed all the experiments with help from R.A.R (preliminary data), R.G., A.G.P., R.O., S.K. (structural biology), S.S. (RNA-seq analysis) and D.J. (peptide arrays). V.C. and L.F. provided the patient samples. I.G. and C.S.H. analyzed the data, prepared the figures and wrote the paper.

## Legends to Figure Supplements and Supplementary Video

**Figure 1 Supplement 1.**
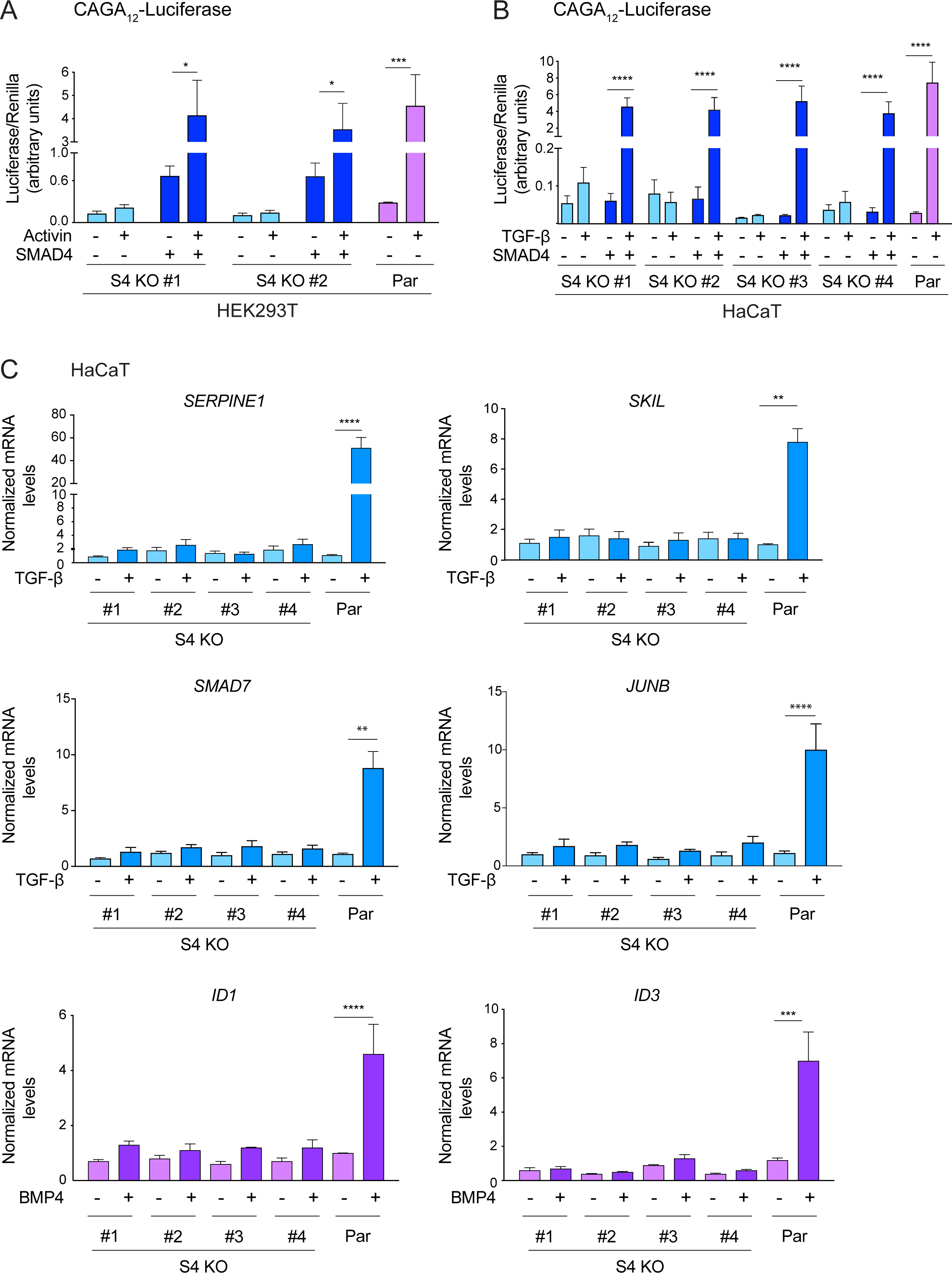
SMAD4 is essential for TGF-β/Activin-induced transcriptional responses. (A and B) HEK293T (A) or HaCaT (B) parental cells and individual clones of SMAD4 knockout cells were transiently transfected with CAGA_12_-Luciferase together with TK-Renilla as an internal control and a plasmid expressing human SMAD4 as indicated (A and B). Cells were incubated with 0.5% FBS-containing media overnight and then treated with 20 ng/ml Activin (A) or 2 ng/ml TGF-β (B) for 8 h. Cell lysates were prepared and Luciferase/Renilla activity was measured. Plotted are the means ± SEM of three independent experiments. The *p* values are from one-way ANOVA with Tukey’s post hoc correction. *, *p*<0.05; ***, *p*<0.001; ****, *p*< 0.0001. (C) HaCaT parental and four independent clones of SMAD4 knockout cells were either untreated or incubated with either 2 ng/ml TGF-β or 20 ng/ml BMP4 for 1 h (*SMAD7, JUNB, ID1* and *ID3*) or 6 h (*SERPINE1* and *SKIL*). Total RNA was extracted and qPCR was used to assess the levels of mRNA for the genes shown. The data are the average of three or four experiments ± SEM. The *p* values are from one-way ANOVA with Tukey’s post hoc correction. **, *p*<0.01; ***, *p*<0.001; ****, *p*< 0.0001. Par, parental; S4 KO, SMAD4 knockout.

**Figure 2 Supplement 1.**
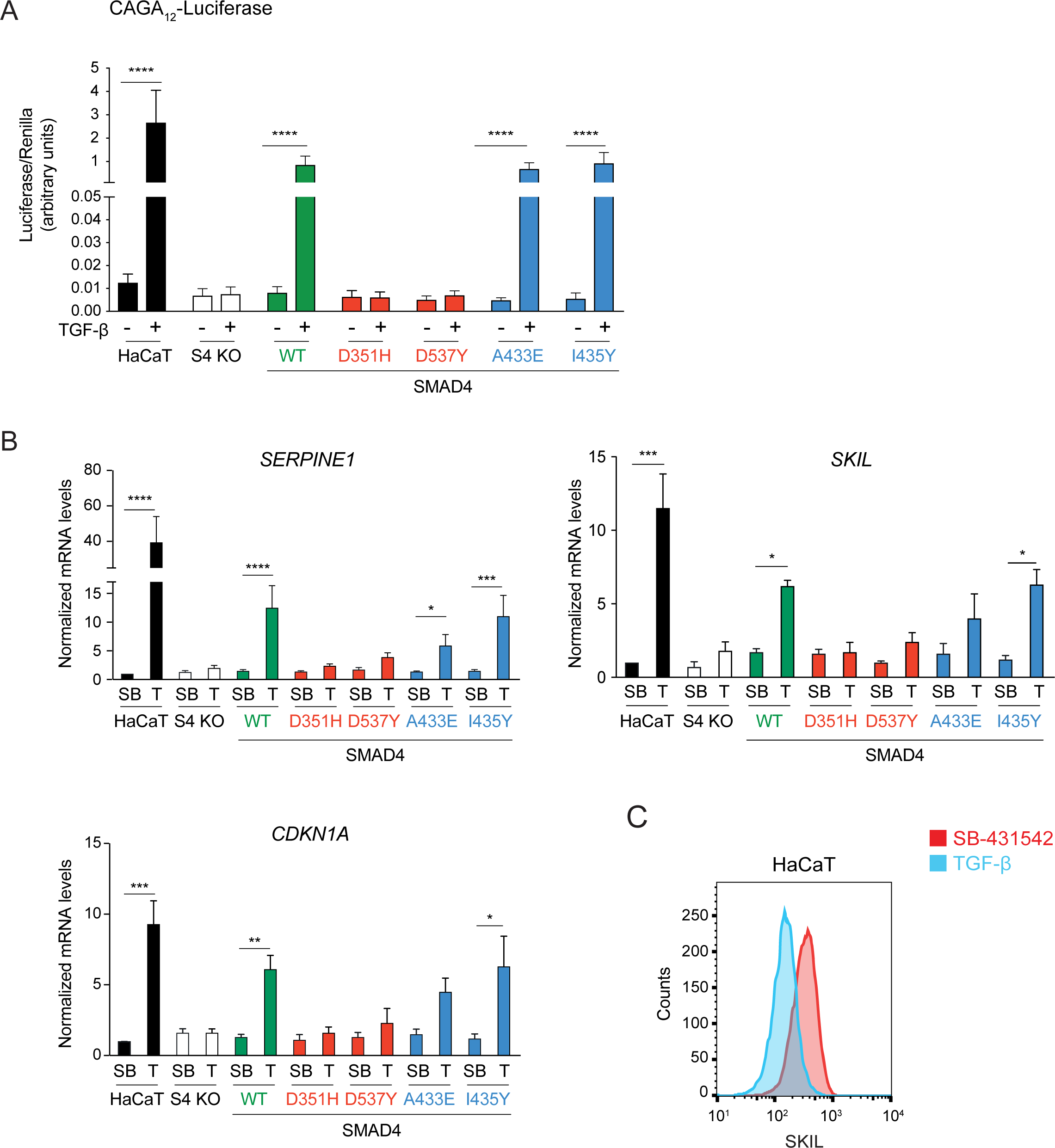
Transcriptional activity of the SMAD4 mutants compared to WT SMAD4. (A) Parental HaCaT, SMAD4 knockout clone 2 stably transfected with EGFP alone (S4 KO), or with EGFP fusions of WT SMAD4 or the four indicated SMAD4 mutants were transiently transfected with CAGA_12_-Luciferase together with TK-Renilla as an internal control. Cells were untreated or treated with 2 ng/ml TGF-β for 8 h. Luciferase/Renilla activity was measured on whole cell lysates. Plotted are the means ± SEM of four independent experiments. The *p* values are from one-way ANOVA with Tukey’s post hoc correction. \*\*\*\**p*< 0.0001. (B) Parental HaCaT and SMAD4 knockout clone 2 cells stably transfected as in (A) were incubated overnight with 10 μM SB-431542, washed out and then retreated with SB-431542 (SB) or with 2 ng/ml TGF-β (T) for 6 h. Total RNA was extracted and qPCR was performed for the genes shown. Plotted are the means ± SEM of four independent experiments. The *p*-values are from two-way ANOVA with Sidak’s post-hoc test. *, *p*<0.05; **, *p*<0.01; ***, *p*<0.001; ****, *p*< 0.0001. (C) Levels of SKIL in parental HaCaT were measured by flow cytometry 1 h after incubation with 10 μM SB-431542 or after treatment for 1 h with 2 ng/ml TGF-β. The panel shows an overlay of the indicated treatment conditions. The red line indicates the SB-431542-treated sample, while the cyan line represents the TGF-β treated sample.

**Figure 4 Supplement 1.**
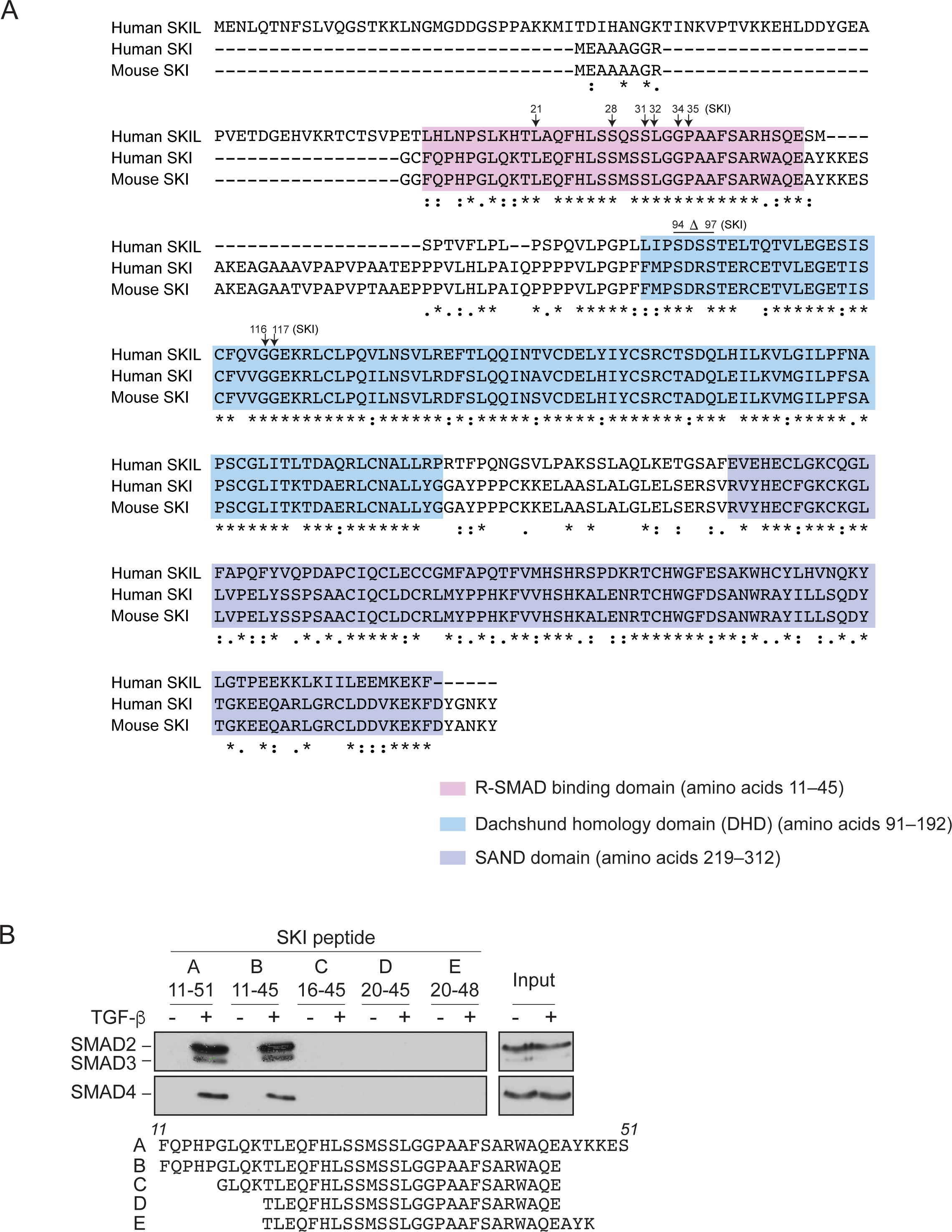
SGS mutations in SKI. (A) Alignment of the first 317 amino acids of human SKI with the corresponding regions of mouse SKI and human SKIL is shown. Key domains are shown: pink corresponds to the R-SMAD -binding domain; blue, DHD domain; purple, SAND domain. SGS mutations are indicated by arrows. (B) SKI peptides corresponding to amino acids 11–51 and truncated versions as shown were analyzed in peptide pulldown assays with whole cell extracts from HaCaT cells treated with or without 2 ng/ml TGF-β. The pulldowns were immunoblotted using the antibodies shown. Inputs are shown on the right.

**Figure 5 Supplement 1.**
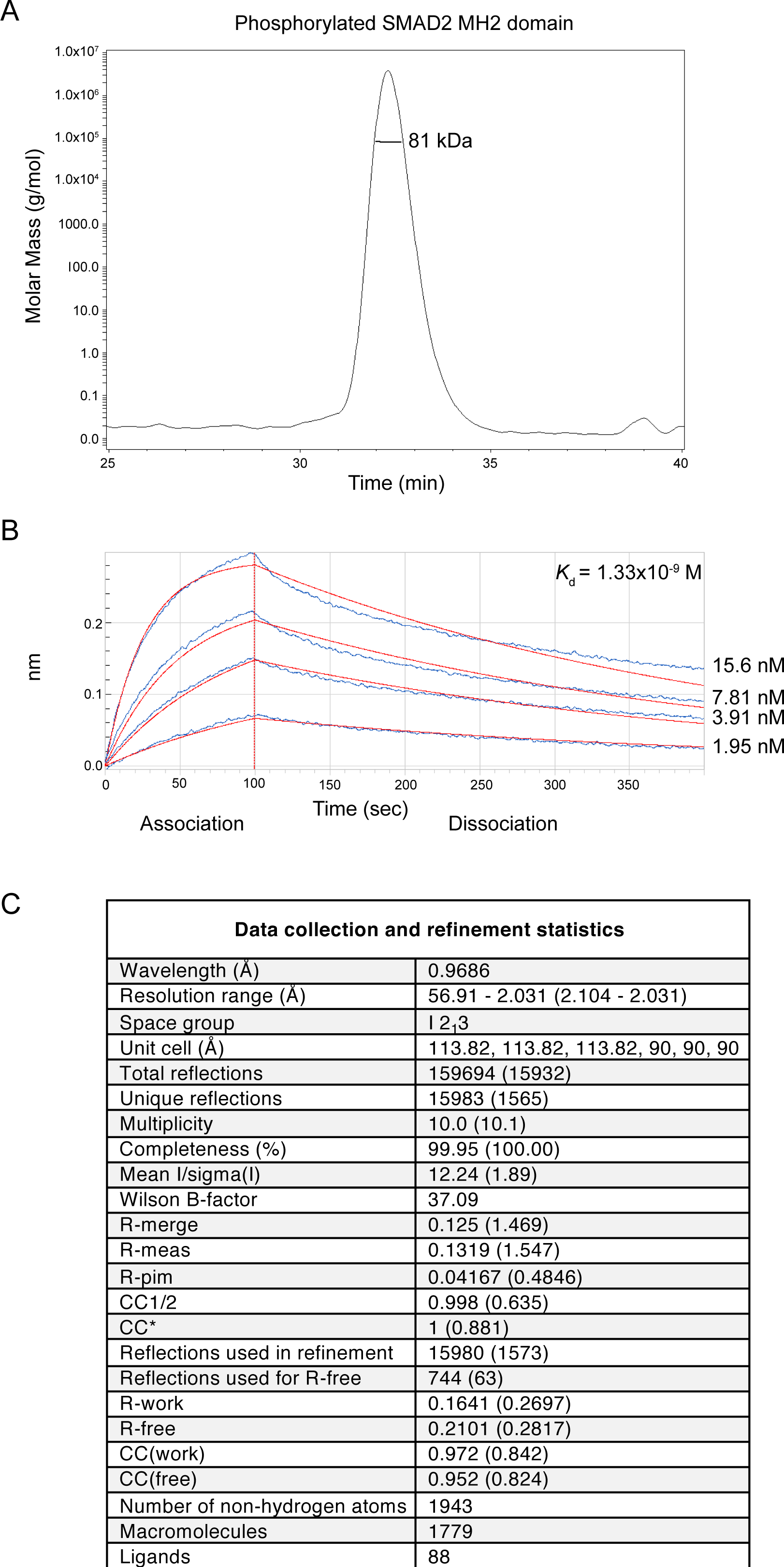
Analysis of the phosphorylated SMAD2 MH2 domain complex used for structural studies. (A) The trimeric arrangement of the phosphorylated SMAD2 MH2 domain was confirmed by SEC-MALLS. The SEC-MALLS chromatogram is shown. The calculated molecular weight was 81 kDa, which was very close to the expected molecular weight of 78.5 kDa. (B) The interaction between the phosphorylated SMAD2 MH2 domain and the SKI peptide (amino acids 11–45) was measured by biolayer interferometry. The biosensors were loaded with biotinylated SKI peptide and incubated with different concentrations of phosphorylated SMAD2 MH2 domain as shown. The calculated *K*_d_ was 1.33×10^−9^ ± 2.12 ×10^−11^ M. (C) Data collection and refinement statistics for the crystal structure of the phosphorylated SMAD2 MH2 domain complex with the N-terminal region of SKI shown in Figure 5A.

**Figure 6 Supplement 1.**
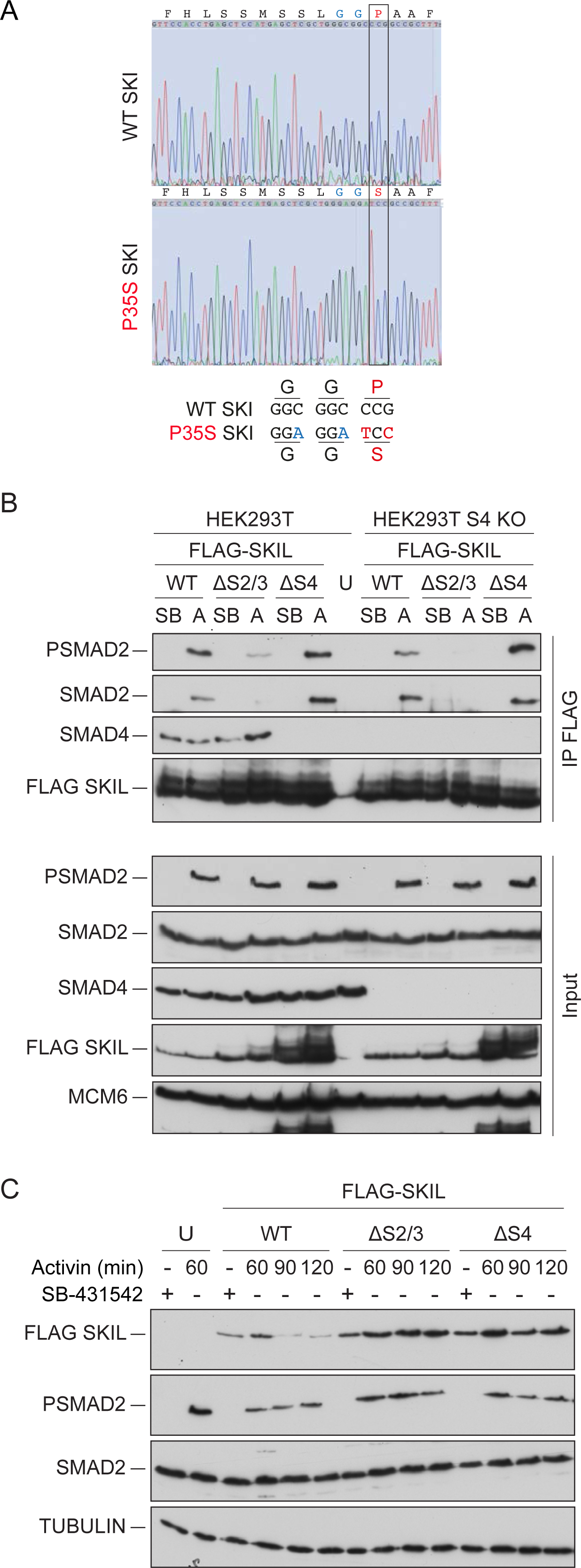
Mutation of the R-SMAD binding domain or SAND domain in SKI/SKIL prevents ligand-induced SKI/SKIL degradation. (A) Characterization of P35S SKI mutation in HEK293T cells compared to parental cells by PCR followed by Sanger sequencing analysis. The black box indicates the change in nucleotides that give rise to the desired mutation. The details of the knocked in changes are given below. (B) HEK293T parental and SMAD4 knockout (S4 KO) cells were transiently transfected with FLAG-SKIL WT or FLAG-SKIL G103V (ΔS2/3) or FLAG-SKIL R314A, T315A, H317A and W318E (ΔS4) as indicated, or left untransfected (U). Cells were incubated overnight with 10 μM SB-431542, then washed out and pre-treated with 25 μM MG-132 for 3 h, followed by incubation with SB-431542 or with 20 ng/ml Activin A for 1 h. Whole cell extracts were immunoprecipitated (IP) with FLAG beads. The IPs were immunoblotted using the antibodies shown. Inputs are shown below. (C) HEK293T cells were untransfected (U) or transfected with the plasmids indicated as in (B). Cells were incubated overnight with 10 μM SB-431542, then washed out and subsequently treated with SB-431542 or with 20 ng/ml Activin A for the times shown. Whole cell extracts were immunoblotted using the antibodies shown. SB, SB-431542; A, Activin A.

**Figure 7 Supplement 1.**
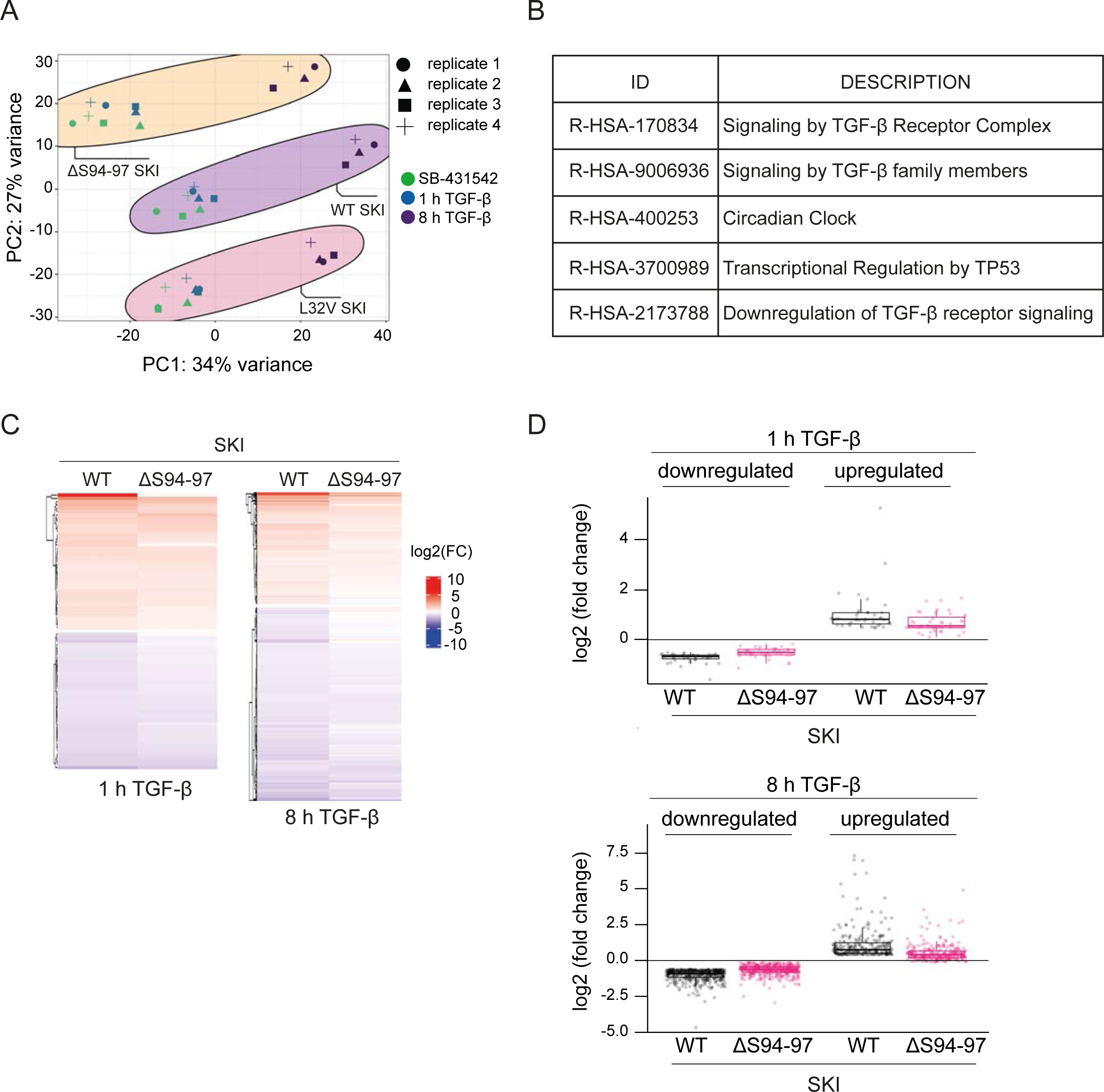
Dermal fibroblasts from SGS patients exhibit an attenuated TGF-β transcriptional response. (A) Principal component analysis (PCA) plot is shown for RNA-seq on normal fibroblasts containing WT SKI, and fibroblasts from SGS patients containing either the L32V SKI mutation or the ΔS94-97 SKI mutation, treated as indicated. PCA measures sample to sample variation using rlog transformed read count of all genes expressed above 1 read in at least one sample. Four replicates are shown for each condition. (B) Enriched Reactome pathways common between the pairwise comparisons of time points (SB-431542-treated versus 1 h TGF-β or 8 h TGF-β) of fibroblasts derived from a healthy subject containing WT SKI. (C) Hierarchically clustered heatmaps of log2FC values (relative to SB-431542 condition) showing the expression of TGF-β-responsive genes in the healthy (WT SKI) and the ΔS94-97 SKI-containing fibroblasts after 1 h and 8 h treatment with TGF-β, analysed by RNA seq. Four biological replicates per condition were analysed. The genes shown are those for which the TGF-β inductions were statistically significant in the healthy fibroblasts, but non-significant in the ΔS94-97 fibroblasts. (D) The same data as in C are presented as box plots.

**Figure 7 Supplement 2.**
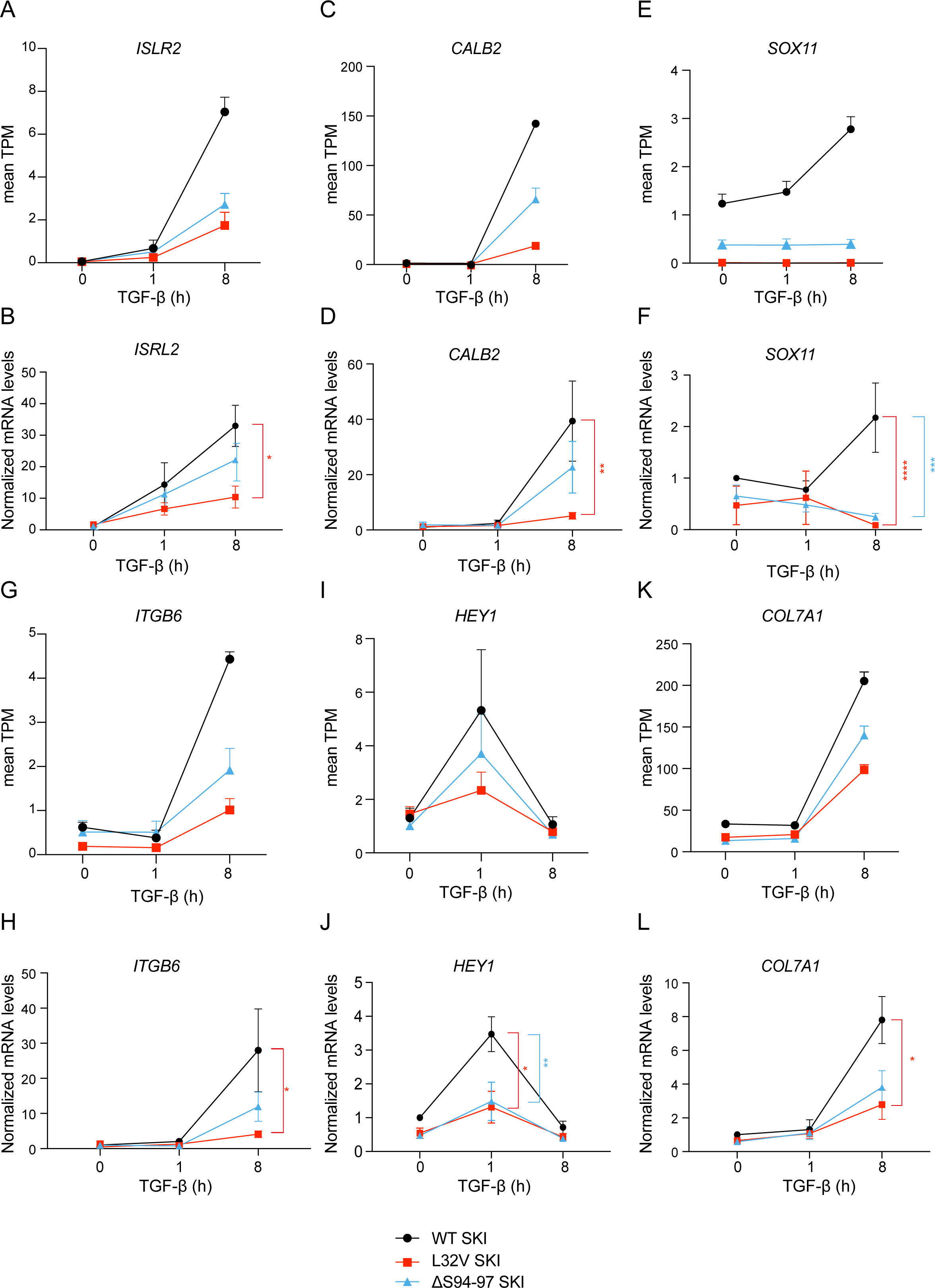
Validation of RNA-seq data by qPCR. (A–L) Healthy dermal fibroblasts (WT SKI) and dermal fibroblasts from SGS patients containing either the L32V SKI mutation or the ΔS94-97 SKI mutation were treated overnight with 10 *µ*M SB-431542, washed out and incubated with either SB-431542 or with 2 ng/ml TGF-β for 1 h and 8 h. Total RNA was extracted and either RNA-seq or qPCR analysis was performed. Transcript levels for a subset of target genes are displayed as plots of the mean transcripts per million (TPM) from the RNA-seq data (A, C, E, G, I, K) or were measured by qPCR and plotted normalized to SB-431542-treated healthy fibroblasts (B, D, F, H, J, L). Plotted are the means and SEM of at least of four independent experiments. The *p*-values are from two-way ANOVA with Tukey’s post-hoc test. *, *p*<0.05; **, *p*<0.01; ***, *p*<0.001; ****, *p*<0.000.

**Supplementary Video 1. Mechanism of SKI binding to phosphorylated SMAD2 MH2 domain**.

Animation of SMAD2 MH2 domain monomer (orange) forming a complex with two other SMAD2 MH2 domain monomers (cyan and olive). Note the movement of Trp448 on helix 5 and Tyr268 on the β1’ strand in the orange monomer upon trimerization. The flipped Trp448 is then in the correct orientation for binding to the SKI peptide (pink).

## Key Resource Table

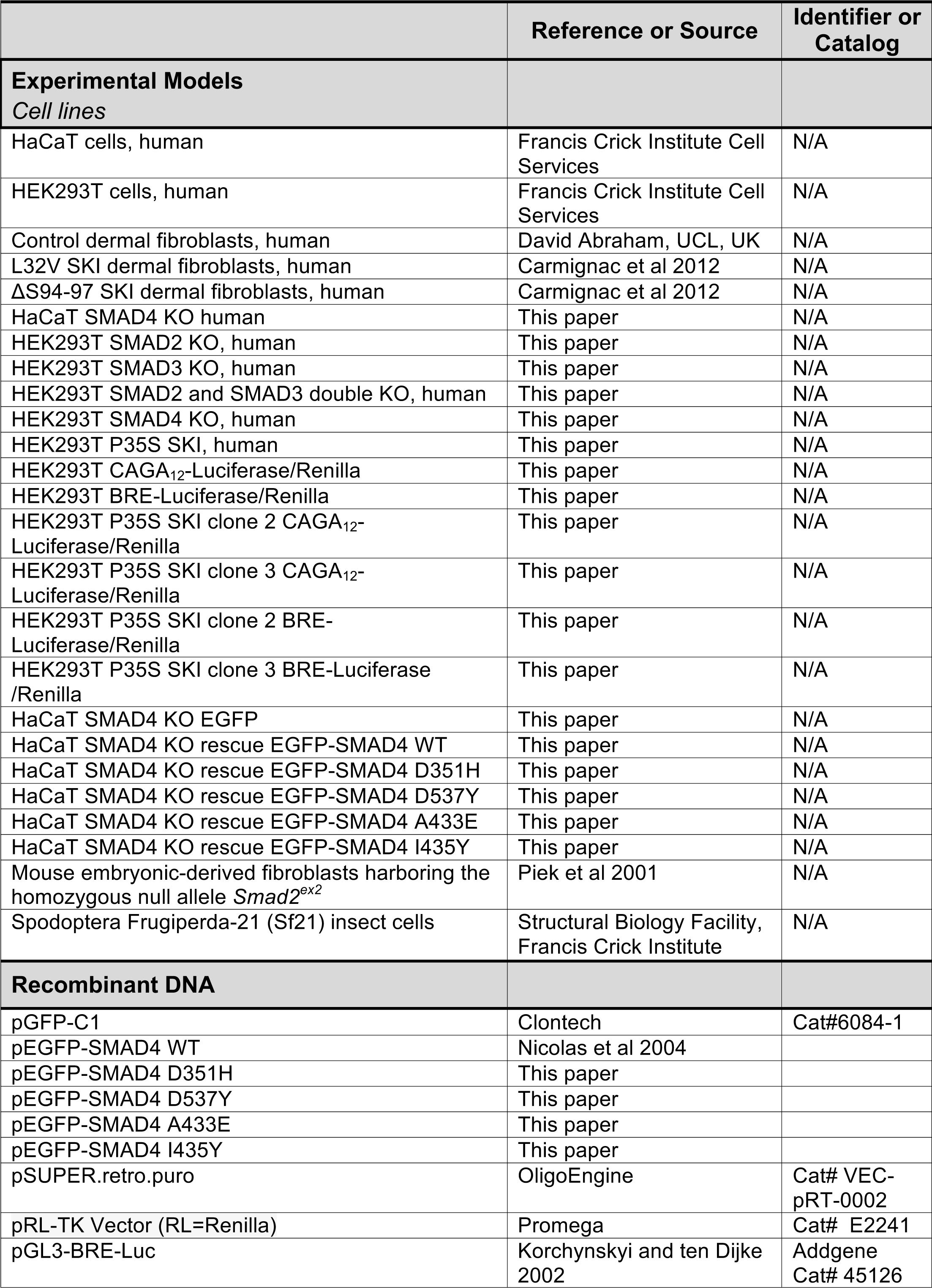
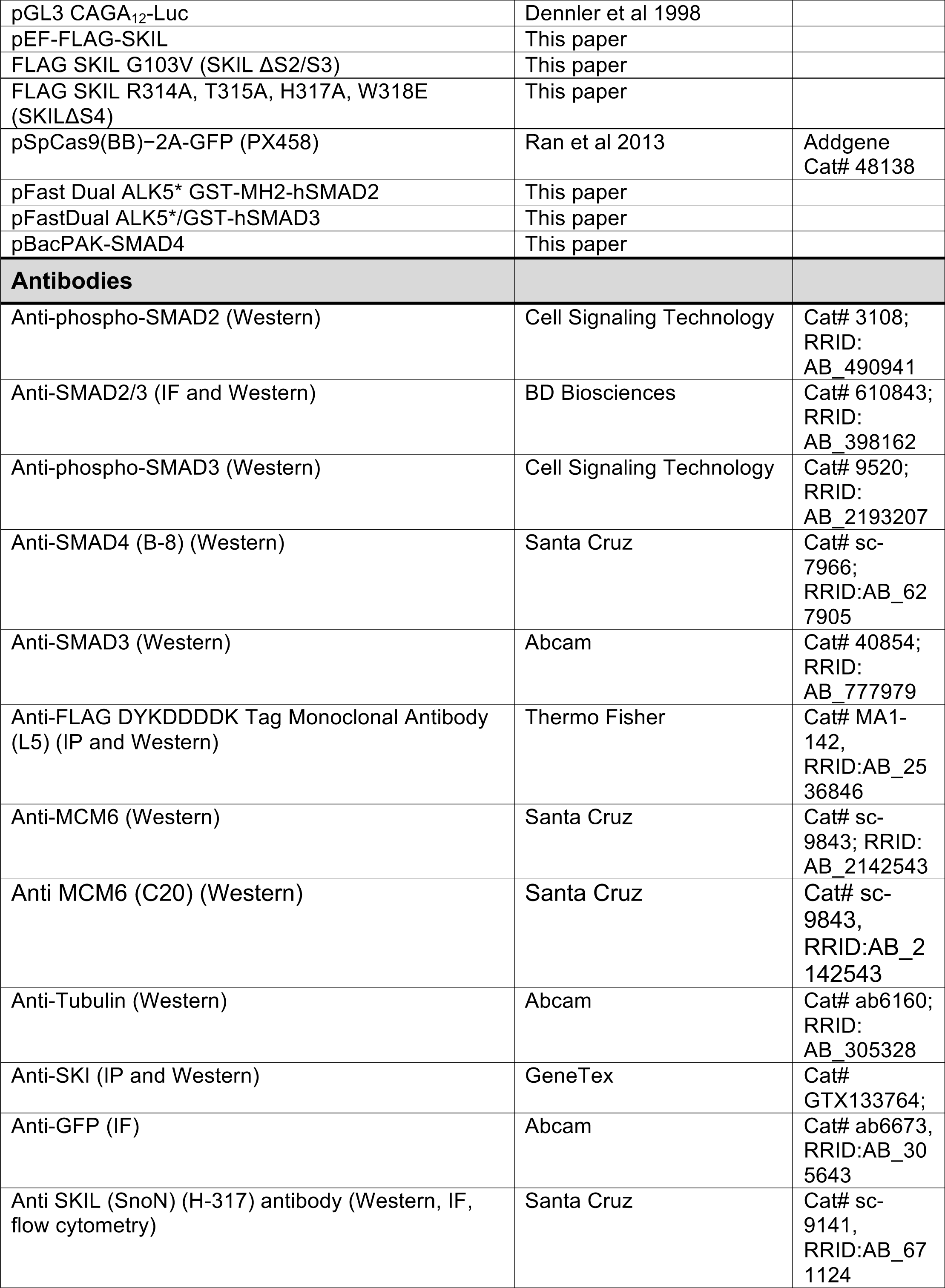
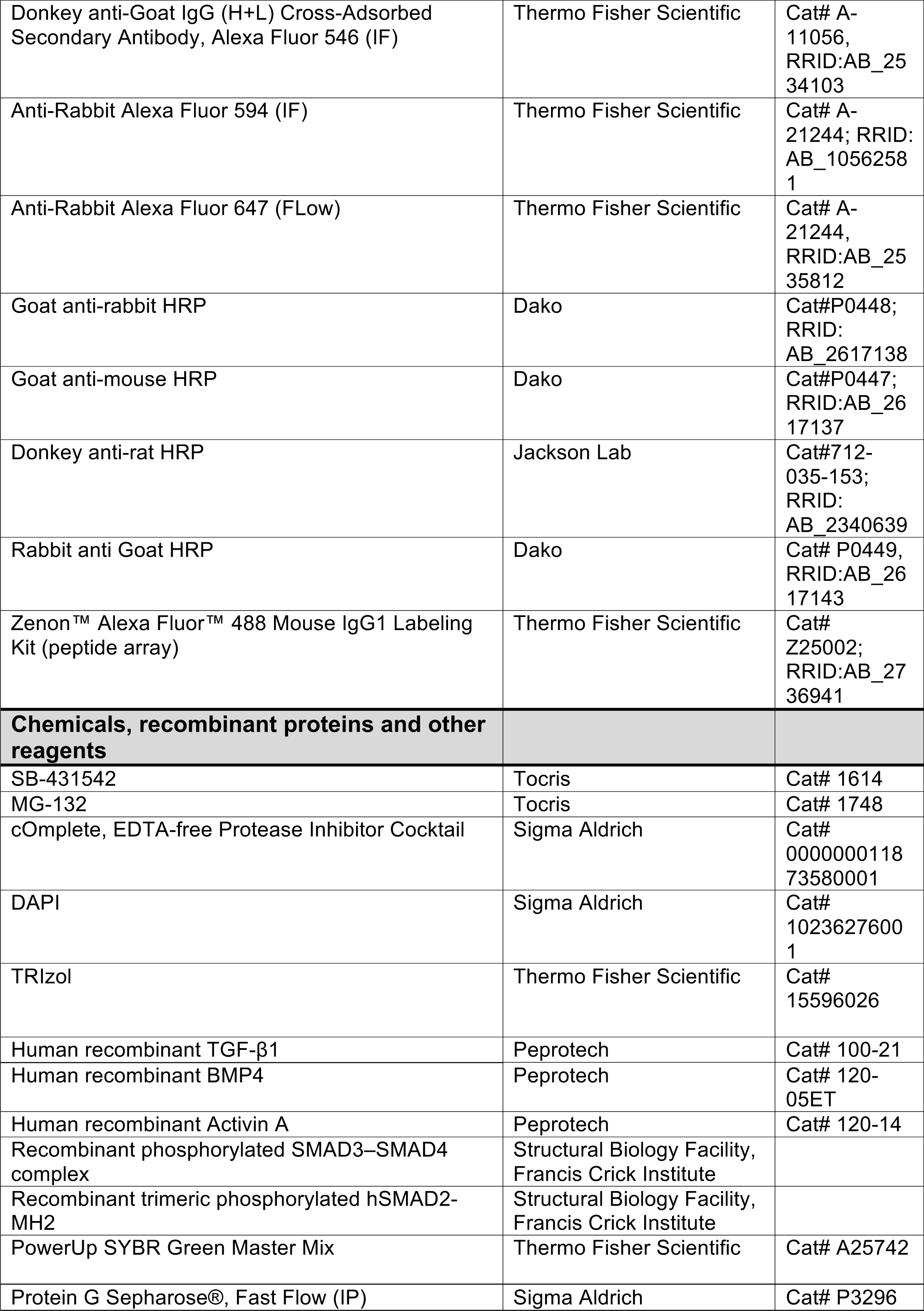
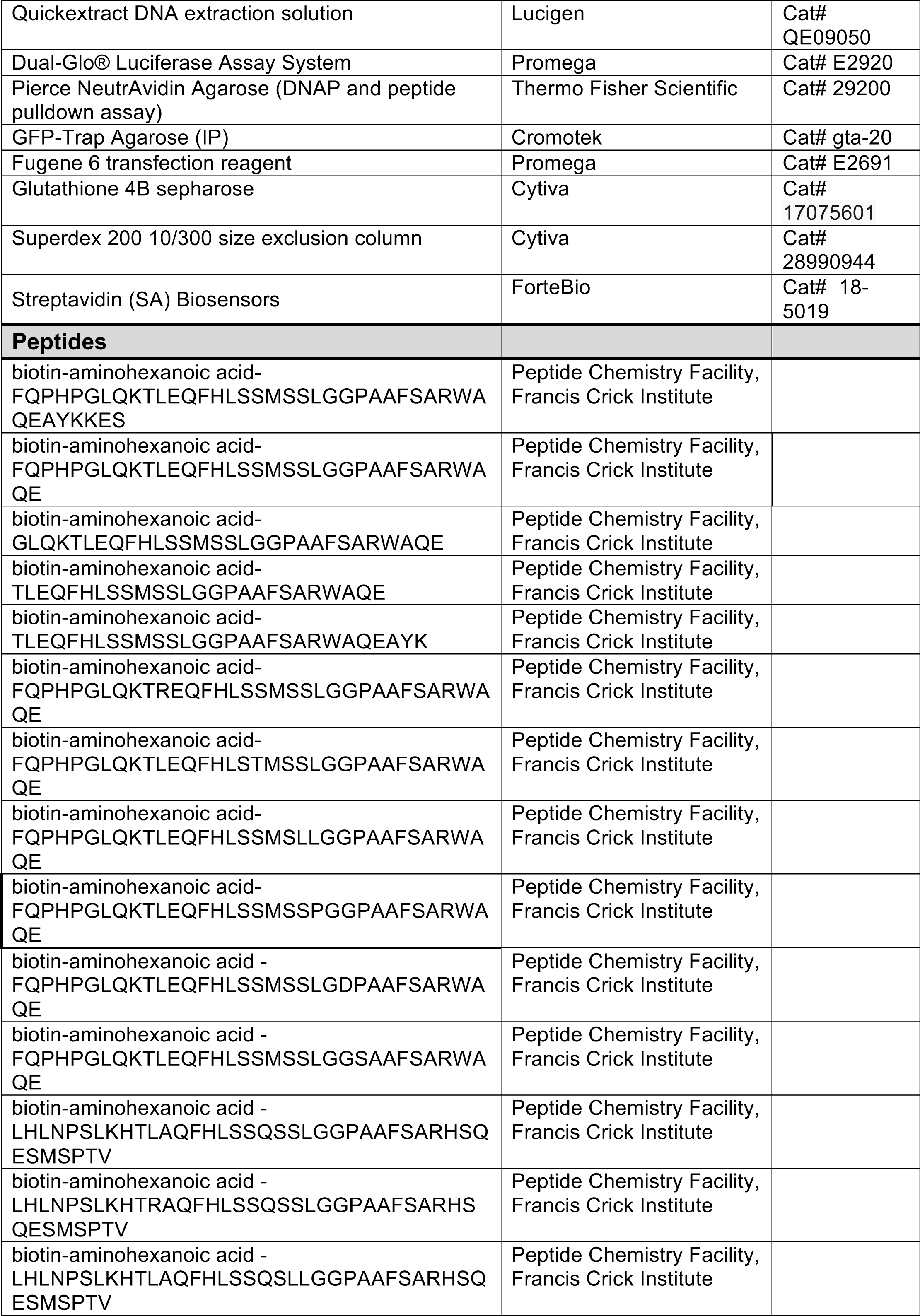
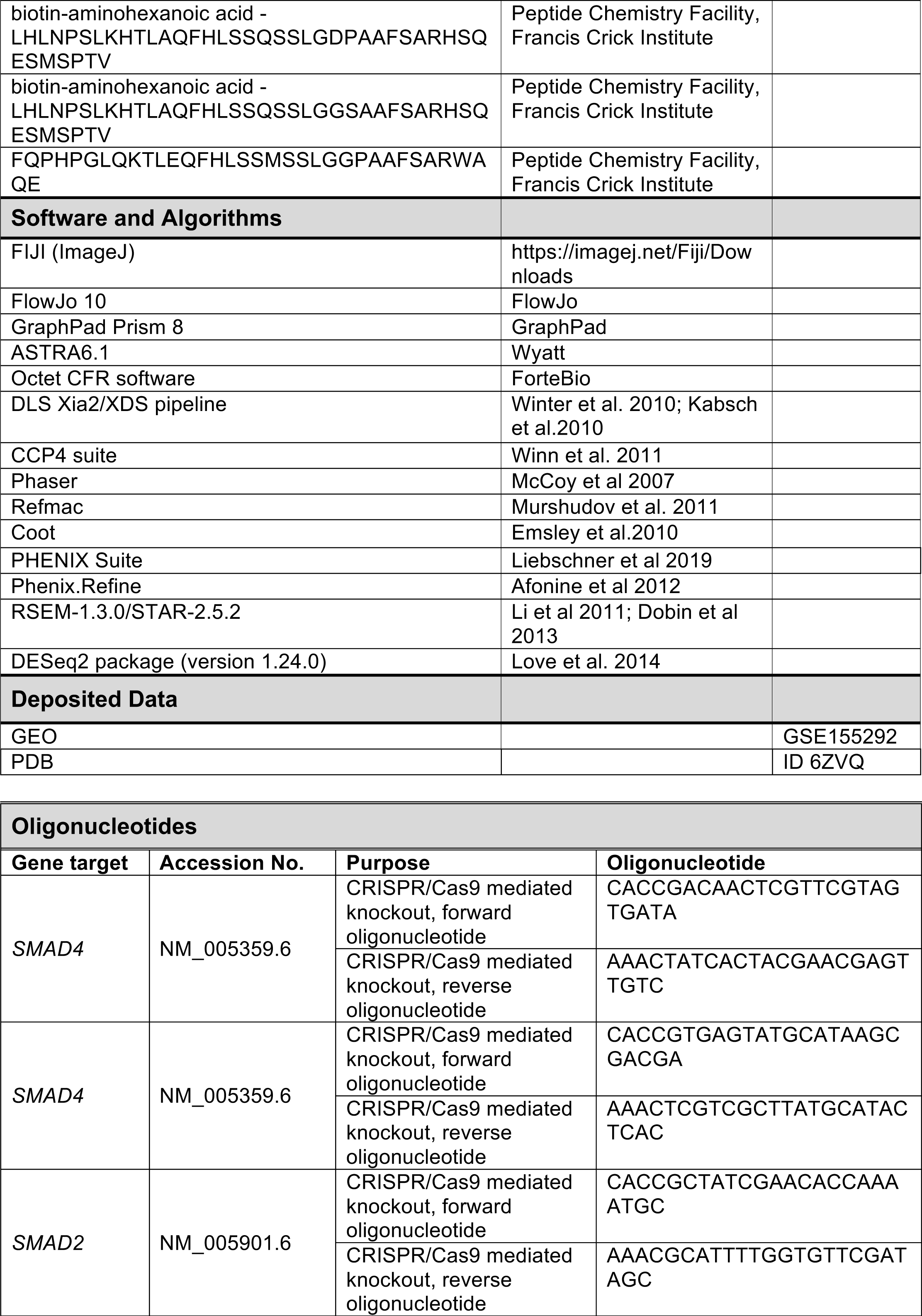
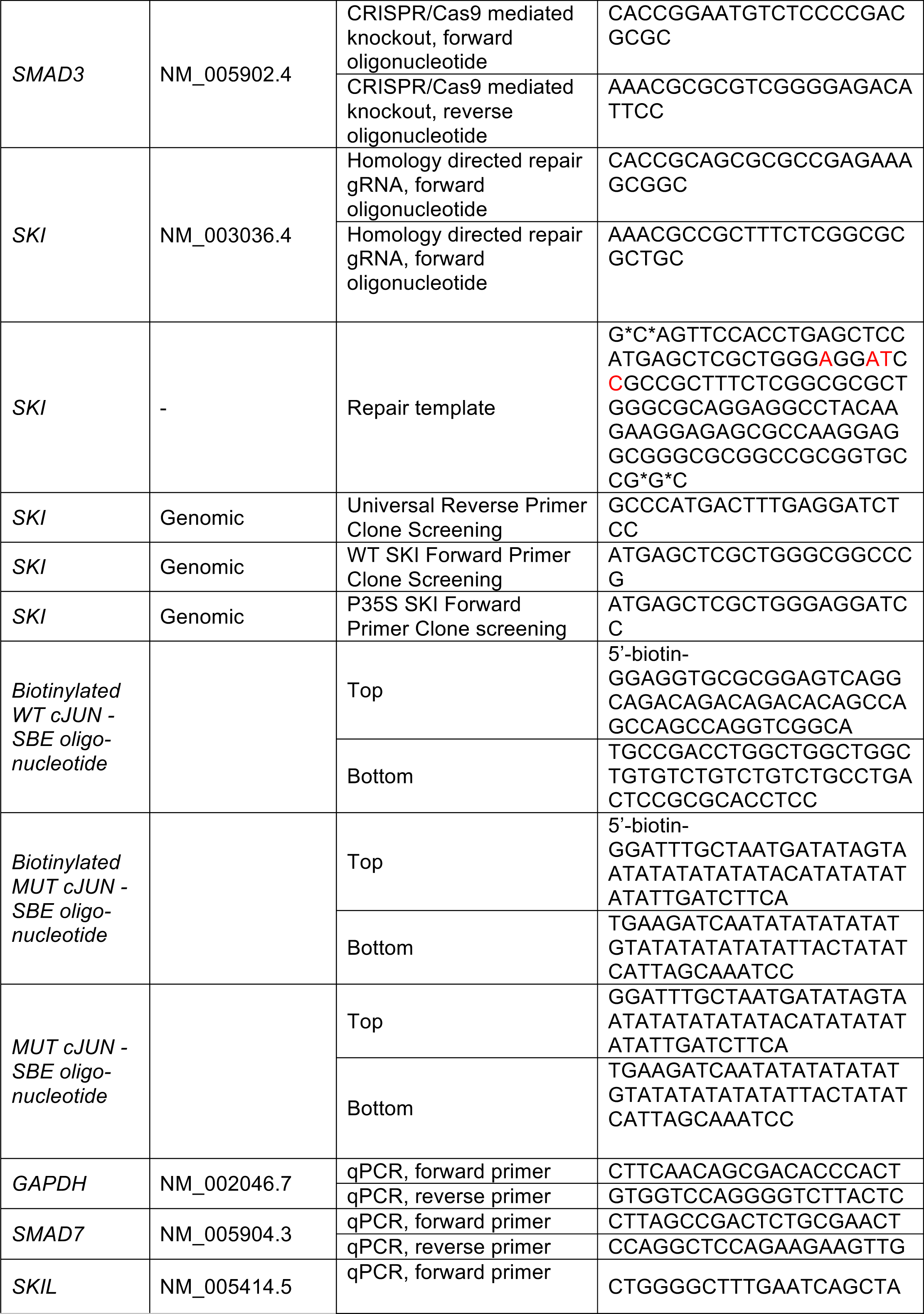
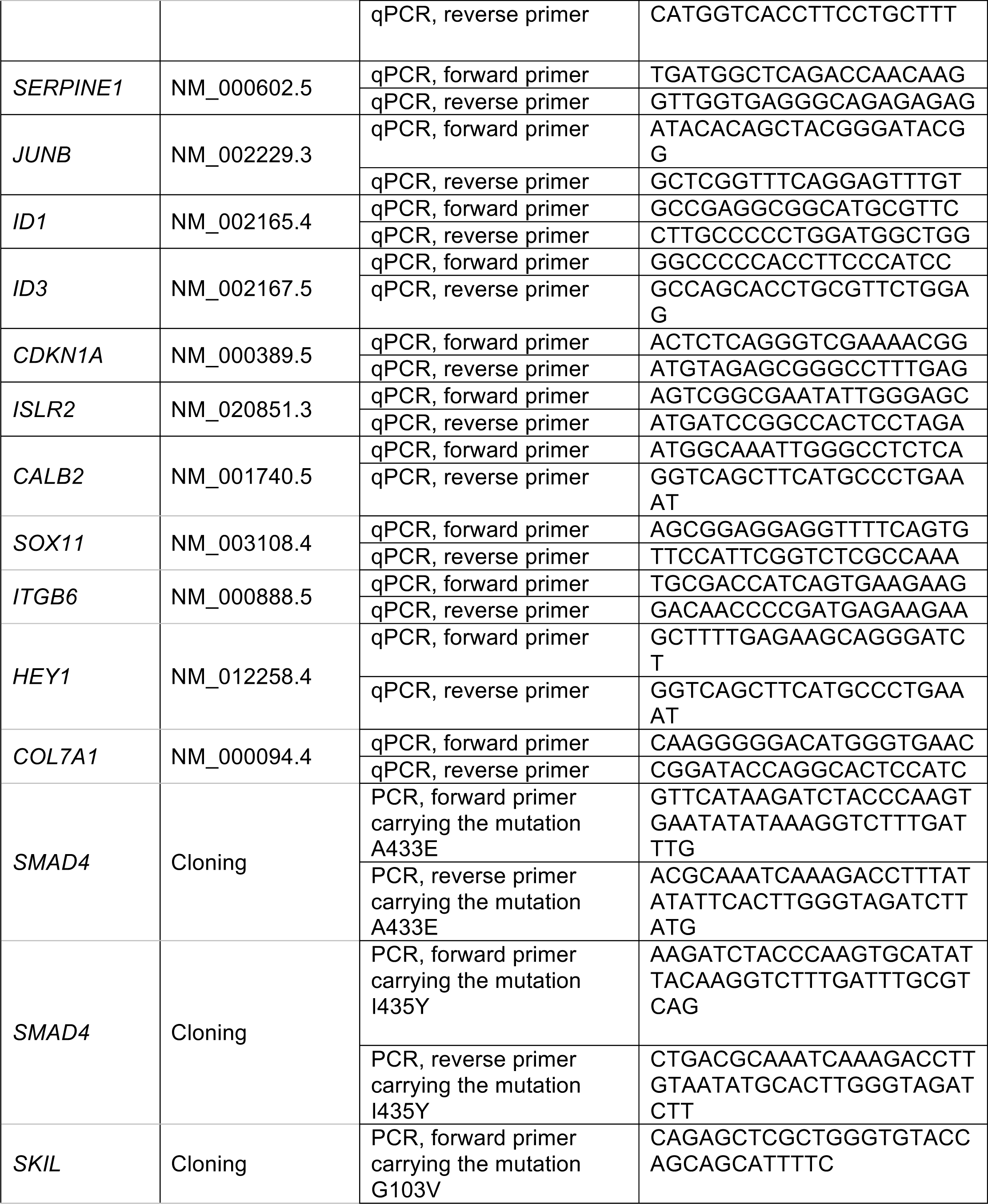

